# Predicting the viability of archaic human hybrids using a mitochondrial proxy

**DOI:** 10.1101/289892

**Authors:** Richard Allen, Hannah Ryan, Brian W. Davis, Charlotte King, Laurent Frantz, Ross Barnett, Anna Linderholm, Liisa Loog, James Haile, Ophélie Lebrasseur, Mark White, Andrew C. Kitchener, William J. Murphy, Greger Larson

**Affiliations:** Palaeogenomics & Bio-Archaeology Research Network, Research Laboratory for Archaeology and the History of Art, University of Oxford, Oxford OX1 3QY, UK; Cancer Genetics Branch, National Human Genome Research Institute, NIH, Bethesda, Maryland 20892, USA; Department of Archaeology, Durham University, Science Site, Durham, UK DH1 3LE; Veterinary Integrative Biosciences, Texas A&M University, College Station TX 77843 USA; Department of Natural Sciences, National Museums Scotland, Chambers Street, Edinburgh EH1 IJF, UK; Department of Anthropology, Texas A&M University, College Station, TX 77843-4352, USA; Department of Zoology, University of Cambridge, Downing Street, Cambridge CB2 3EJ, United Kingdom; School of Biological and Chemical Sciences, Queen Mary University of London, Mile End Road, London E1 4NS, UK; Institute of Geography, School of Geosciences, University of Edinburgh, Drummond Street, Edinburgh EH9 3PX, UK; Department of Anatomy, University of Otago, Great King Street, Dunedin 9016, NZ

## Abstract

Ancient DNA evidence has confirmed hybridization between humans and Neanderthals and revealed a complex pattern of admixture between hominin lineages. Many segments of the modern human genome are devoid of Neanderthal ancestry, however, and this non-random distribution has raised questions regarding the frequency and success of hybridisation between ancient human lineages. Here, we examine the hypothesis that hominin hybrid offspring suffered a reduction in fertility by comparing patterns of sequence divergence of mitochondrial and nuclear DNA from numerous hybridising pairs of mammals. Our results reveal a threshold separating species pairs whose divergence values fall within two categories: those whose hybrid offspring can successfully reproduce without backcrossing with their parent species, and those whose hybrid offspring cannot. Using this framework, we predict that the potential hybrid offspring of Neanderthals, Denisovans, the ancient individuals from the Sima de los Huesos and anatomically modern humans would not have suffered a reduction in fertility.

## Introduction

Though numerous interspecies hybrids have been recorded, predicting their viability and relative fertility has proved difficult. For example, the absence of Neanderthal mitochondrial genomes in the modern human population led some to suggest that anatomically modern humans (AMH) and Neanderthals did not hybridise^1-3^. Recent analyses of whole genome sequences derived from ancient individuals, however, have demonstrated that archaic hominins including Neanderthals and Denisovans did in fact hybridise with AMH^4-6^. For many anthropologists, this result corroborated previous conclusions (based on morphological grounds) that Neanderthals and AMH could have produced fertile offspring (*e.g.*^7^).

A recent paper, however, conflated the observation of genetic deserts devoid of Neanderthal ancestry on modern human autosomes, and especially the X chromosome, with reproductive incompatibilities between humans and Neanderthal^8^. Though this study also suggested that demographic factors likely played a role in the formation of these deserts, two studies^9,10^ concluded that most selection against Neanderthal DNA in genic regions was not due to reproductive incompatibilities, but instead was the result of the Neanderthals’ high genetic load. In addition, the initially high proportion of Neanderthal admixture^11^ suggests that early male and female hybrids of AMH and Neanderthals were normally fertile, and that any Neanderthal variants that may have interfered with fertility within an AMH genetic background were likely confined to a few genetic loci. Lastly, multiple factors including ecological drivers can lead to genomic deficits of introgression within hybrid offspring that are otherwise fertile^12^, suggesting the existence of genetic deserts may not imply reproductive incompatibility.

More generally, assessing whether any two species are capable of producing viable or fertile offspring is difficult in the absence of nuclear genomic data, captive experiments or field data. In addition, there has been no robust, quantitative biological measure of the expected fitness of hybrid offspring, and thus no rigorous method to ascertain the specific viability of archaic human hybrids, or the potential for hybridisation between more ancient hominin lineages. Determining whether distinct hominin lineages were capable of producing fertile offspring is important not only for understanding the patterns of hybrid ancestry within modern humans, but also for ascertaining the degree to which reproductive isolation (as a result of either cultural or biological phenomena) existed between AMH and other hominin lineages.

Though numerous studies have speculated about the potential for hominin lineages to produce viable hybrids, few have made use of the numerous pairs of mammalian species known to produce hybrid offspring along a continuum of male (and to a lesser extent, female) fertility. Based on the qualitative correlation that the fertility of hybrids generally decreases with divergence time, two recent studies^13,14^ speculated that, given their relatively recent temporal divergence, AMH and Neanderthals could have retained the ability to produce fertile offspring. These studies could not rule out the fact that the male hybrids may have possessed lower fertility as a result of Haldane’s Rule which states that if one sex is rare, absent or sterile among hybrid offspring, it is the heterogametic sex.

In order to obtain a quantitative measure of whether two lineages can interbreed, we developed a framework based on the correlation between mitochondrial genetic distance between mammalian species pairs, and the manifestation of hybrid male sterility. Some previous studies have questioned this approach and stated that measures of species divergence are not necessarily reliable predictors of hybrid sterility^15,16^. Other studies have reported that genetic divergence values do correlate with species boundaries^17^ and time to speciation. A more specific recent study of damselflies demonstrated a strong correlation between the genetic distances separating two species and their relative reproductive isolation^18^.

Hybrid fertility exists along a continuum. For the sake of simplicity and ease of distinction, we first explicitly defined two dichotomous categories amongst hybrid offspring. Category 1 contains eight terrestrial mammalian species pairs that are capable of producing F1 offspring that can reproduce without backcrossing with a parent species (even if there are observed asymmetries in gene flow and variation in male fertility amongst the hybrids) (Table S1). Category 2 consists of nine pairs of species that can produce viable F1 offspring, but these hybrids either cannot reproduce without backcrossing with a parent species, or are completely infertile (Table S1). In order to determine the categorical assignment of each species pair, we relied upon empirical evidence derived from experimental studies of F1 hybrid fertility (Table S1). We also compiled 22 additional species pairs that are known to produce viable offspring, but for which there was insufficient evidence to confidently assign them into either category (Table S2, Fig. S2).

We then aligned sequences of both the cytochrome b gene (*CYTB*) and mitochondrial genomes (excluding the control region) from multiple individuals per species and confirmed (by matching the phylogenies to nuclear species trees) that the selected mitochondrial sequences for each taxon pair were neither mislabelled, nor nuclear copies of mitochondrial genes, nor derived from hybrid populations. From the sequence alignments we calculated average pairwise genetic distances between each species pair using both Hamming (raw) distances and evolutionary models. In addition, we calculated genetic distances between ten primate species pairs using four nuclear loci (*CHRNA1, GHR, ZFX, ZFY*)^19^ (Fig. S3).

## Results and Discussion

The calculated divergence estimates using *CYTB* revealed clear thresholds distinguishing the two categories (Fig 1, Fig. S1). More specifically, none of the eight Category 1 pairs that were capable of producing fully fertile offspring possessed *CYTB* sequences with raw average pairwise divergence values greater than 7.60%. Other than two vole species that possessed a divergence value of 7.54%, all other Category 2 pairs possessed divergence values greater than 7.73%. The existence of a genetic distance threshold also held true for the mitogenomes (Fig. S3). In addition, the male and female hybrid offspring of the two most divergent pairs were shown to be completely infertile corroborating previous studies^20-22^ showing that along the continuum of speciation, infertility evolves prior to inviability. Lastly, a Mann-Whitney U test showed significantly lower genetic divergence values of species pairs in Category 1 relative to those in Category 2 (p < 0.0001).

**Figure 1.**
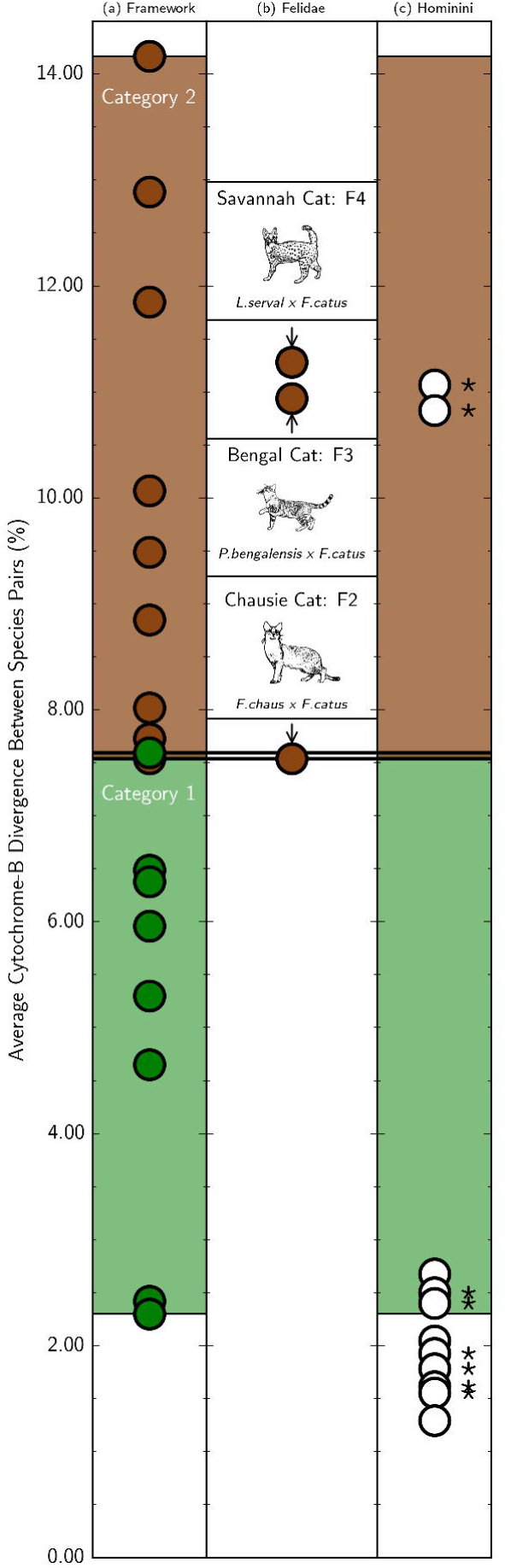
A depiction of the correlation between *CYTB* divergence between terrestrial mammalian species pairs and the relative fertility of their hybrid offspring. In column A, the green circles represent species capable of producing fully fertile F1 offspring which can reproduce independently of their parent species (Category 1). Brown circles represent species pairs whose F1 offspring either require backcrossing with a parent species or are completely infertile (Category 2). The lighter green and brown shaded regions represent the range of divergence values of the two categories and, given their general lack of overlap, can be used to determine the relative fertility of hybrid offspring. Column B depicts the divergence between three wild felid species and domestic cats, as well as the minimum number of generations of backcrosses with domestic cats before full fertility of the hybrid is restored. The white circles in Column C depict the divergence between three ancient hominins and AMH, as well as the distances between AMH and chimpanzees and bonobos (in Category 2). The asterisks represent those pairs that include modern samples of AMH. The lack of an asterisk signifies that only sequences derived from archaeological AMH were used to compute the divergence values. All hominin pairs are positioned at the lowest (most fertile) end of Category 1, suggesting that their hybrid offspring are likely to have been fully fertile. Additional detail regarding the specific species pairs are listed in Fig. S1 and Table S1.

Importantly, the two categories of fertility defined here are not strictly linked with gene flow. For instance, though both male and female hybrid offspring of all species pairs in Category 1 can reproduce without requiring a backcross with a parent species gene flow asymmetries have been demonstrated in virtually all Category 1 pairs including house mice (*Mus musculus musculus* x *Mus musculus domesticus*)^23^, and brown (*Ursus arctos*) and polar bears (*Ursus maritimus*)^12^ (Table S1). In addition, gene flow has also been demonstrated between Category 2 species (including *Mus musculus* and *M. spretus*^*24*^) whose F1 hybrid offspring cannot produce viable F2s. Since both fertility and the potential for gene flow vary along a continuum, it is interesting to note that the divergence values associated with the two fertility categories defined here do not overlap more substantially. As a result, our results suggest that pairwise mitochondrial genetic distance values can be used to make predictions of male hybrid sterility, at least in terrestrial mammals.

Though the lack of available sequences for nuclear loci (relative to mitochondrial loci) limited our ability to test whether nuclear genes produced the same general pattern as the mitochondrial loci across all our species pairs, we were able to identify four nuclear genes: Zinc finger Y-chromosomal protein (ZFY), Zinc finger X-chromosomal protein (ZFX), Growth Hormone Receptor (GHR), Cholinergic Receptor Nicotinic Alpha (CHRNA1), that had been sequenced in 10 of the primate pairs (Fig. S3) that were known to produce viable hybrid offspring^19^. We generated pairwise distances for each of these genes using the same methodology employed in the mitochondrial analyses. We then assigned each species pair to Category 1 or Category 2 based upon their CYTB divergence values within the original framework. In each case, though the order of the taxa based upon pairwise divergence values varied relative to the pattern generated using *CYTB* (owing to the significantly smaller divergence values in nuclear loci), for each locus the limited overlap in divergence values associated with the two categories was consistent with the mitochondrial assessment (Fig. S3).

In order to further substantiate both this correlation and the robustness of *CYTB* as a proxy for hybrid fertility, we tested the utility of this system for predicting fertility in a well-known hybrid system. To create hybrid pets, cat breeders have crossed domestic cats (*Felis catus*) with three species of wild felid: Jungle cats (*F. chaus)*, Leopard cats (*Prionailurus bengalensis*), and Servals (*Leptailurus serval*)^*25*^. In all cases, the first generation male hybrids are sterile. To regain fertility while maintaining some wild cat characteristics, breeders must backcross the F1 female offspring with male domestic cats to establish a breeding population of pets^*25*^. Given that backcrossing is required for all three crosses, our framework would firstly predict that the *CYTB* distances between all three pairs should be close to or greater than ∼7.5% and that they should all fall into the Category 2 range. Secondly, the pairs with greater genetic distances should require more numerous backcrosses with domestic cats (halving the wild cat ancestry with each subsequent generation) before full fertility is restored and a breeding pet population is established.

Both of these predictions are borne out by the data (Fig 1, Table S3). All three pairs possess *CYTB* distances greater than or equal to 7.54% and the increasing molecular distances between the pairs correlate with an increase in the number of required backcross generations to regain fertility. Specifically, distances between Jungle cats, Leopard cats, Servals and domestic cats (7.54%, 10.94%, and 11.28% respectively) are consistent with both the observed minimum (2, 3, and 4 respectively) and average (3, 4, and 5, respectively) number of backcrosses with domestic cats required for hybrid males to acquire fertility^25^. These results are also consistent with an early hybrid experiment using guinea pigs in which hybrids between *Cavia fulgida* and *C. porcellus* (8.02% *CYTB* divergence) were able to regain male fertility after the third generation of backcrosses^26^ (Table S1).

Recent accidental hybrids in zoos also confirm the predictive power of this proxy. In 2006, the Copenhagen Zoo placed a domestic sow (*Sus scrofa domesticus*) in a pen with a male babirusa (*Babyrousa celebensis*) with the expectation that the two species were sufficiently evolutionarily divergent that they would be incapable of producing offspring. The *CYTB* divergence between the two species (12.89%), however, falls within the Category 2 range. Months later, five infertile piglets were born and though two died from maternally induced trauma, the other three (two males, and one female), all survived^27^ (Table S1, Fig. S1). Historically, zoos have often accidentally produced hybrids offspring between distantly related species (Table S2), though the relative fertility of the F1s was rarely established.

Having approximated the *CYTB* threshold values within the framework, we then predicted the relative fertility of hybrids between pairs of other hominin lineages. To do so, we calculated the average pairwise divergence in *CYTB* sequences between AMH and three extinct hominin lineages: Neanderthals, Denisovans, and the ancient population from the Sima de los Huesos cave in Spain^28^. To avoid overestimating the genetic divergence that results from comparing modern and extinct populations, we also generated distance values using the CYTB sequences derived solely from ancient AMH found in archaeological contexts.

The divergence values for each pairing between three *Homo* groups (Sima de los Huesos, Neanderthals, and AMH) all possess divergence values at the bottom of the Category 1 range (Fig 1, Fig. S1, Table S1). Interestingly, the divergence values of Denisovan-Neanderthal and Denisovan-AMH are the largest of the *Homo* pairings, and are consistent with the suggestion that Denisovans possess a mitochondrial lineage that may have introgressed from another source population^29^.

In order to assess hybrid fertility within the hominin lineage more deeply, we calculated the pairwise *CYTB* divergence between humans and our closest living relatives: Chimpanzees (*Pan troglodytes*) and bonobos (*P. paniscus*). Female chimpanzees inseminated with human sperm during a Soviet experiment in the 1920s failed to produce any offspring, and the reverse experiment did not progress beyond the planning stage^30^. Recent molecular clock assessments have suggested that AMH and chimps diverged ∼5-6 Mya^31^, well beyond both the two million year threshold purported by other studies as the upper limit to hybrid fertility^14^, and the average time to speciation^32^.

Interestingly, our analysis places the divergence values between AMH and chimps, and AMH and bonobos within Category 2 (Fig 1, Fig S1). Six other species pairs (including two of the wild cat and domestic cat pairs) possess divergence values equivalent to or greater than AMH, chimps, and bonobos (Table S1, Fig. S1). In all six cases, however, each of the pairs possesses an identical number of chromosomes. Though karyotype is not a strict barrier to the production of hybrids (Table S1), AMH possess two fewer chromosomes than chimps and bonobos (Table S1).

## Conclusions

When placed within the context of other mammalian species, ancient hominin lineages were likely not sufficiently divergent from each other to expect a biological impediment to the generation of fertile offspring. Although some studies have inferred a low level of introgression^33^, the relative ease of hybridisation between hominin lineages is demonstrated by the fact that archaic populations interbred with AMH on at least four occasions^34^. The absence of a biological barrier shifts the focus toward cultural considerations and demographic factors to explain the lack of Neanderthal and Denisovan genomic contribution on the AMH X chromosome, and the lack of mitochondrial DNA from either species within AMH. Palaeoanthropologists disagree as to what differences in culture, behaviour and cognition existed between AMH and Neanderthals (e.g.^35,36^). Though these factors may have altered the sexual symmetry of the hybrids, they appear not to have precluded introgression in either direction^37^.

This study demonstrates the power of genetic distances as a proxy to predict the relative sterility of hybrids resulting from matings between mammalian species. Hybrid sterility is both fluid and governed by numerous mechanisms at multiple levels of biological organization. In addition, gene flow between species whose hybrids require backcrossing is possible (Table S1), as are asymmetries in gene flow in species pairs whose F1 offspring can produce fertile F2s. We do not claim that *CYTB* plays a causative role in hybrid fertility (though a recent study proposed that speciation may be mediated by mitonuclear interactions^38^), and the use of genetic divergence values as a proxy cannot be perfectly predictive. For example, under the Dobzhansky-Müller model, incompatibility can arise from as few as two mutations in each of the admixing populations irrespective of time since divergence, meaning that it would be possible for closely related populations to be incapable of generating fertile hybrids^39^, though no such examples have yet been described.

The value of any proxy, however, is determined by both its predictive power and the ease of generating the proxy data. Publicly available mtDNA sequences from thousands of mammalian taxa already exist and calculating pairwise divergence values is inexpensive, simple and fast. As a result, *CYTB* specifically (and genomic distances more generally) have substantial value as a means to predict the potential for any two mammalian species to produce viable offspring, and the relative degree of the hybrid’s sterility. The discovery of additional extinct hominin populations that survived into the last 250,000 years, including *H. floresiensis* and *H. naledi*, has raised interest in understanding the limits to fertility and hybridization between extinct and extant *Homo* spp. If and when mitochondrial genomes from these samples can be obtained, the approach described here may provide an answer, even if nuclear genomic data are not obtainable.

Lastly, establishing which species pairs violate the predictions of the framework will lead to a better understanding of the process of speciation and the biological and sociocultural mechanisms responsible for hybrid sterility. Our framework can also be applied more generally in contexts such as conservation biology, where predicting the consequences of biological hybridization is key to policy considerations.

## Materials and Methods

### Assessment of Hybrid Fertility and Rationale of Assignment into Categories

In order to ascertain if there was a relationship between genetic divergence and the fertility of hybrid offspring between species, we first collected published examples of species pairs that were capable of producing live offspring. We then split the hybrid pairings into two categories. Into Category 1 we placed eight species pairs that are capable of producing fertile F1 offspring of both sexes that can reproduce without backcrossing with a parent species. Though within these examples there have been observed asymmetries in gene flow and variation in fertility amongst the hybrids, none of these examples requires (one or more generations of) backcrossing with a parent species for their offspring to regain fertility. More specifically, for an example to be listed in this category, we required evidence of fertility in both male and female F1 hybrids. The evidence and rationale for placing each of these pairs into Category 1 is listed in Table S1.

The hybrid offspring of all of nine pairs of species in Category 2 either require one or more generations of female hybrids backcrossing with the male of a parent species to produce fertile offspring, or are completely infertile. For these pairs, we obtained evidence demonstrating no successful F2s from F1 hybrid couplings, an inability to produce offspring other than by backcrossing to a parent species, or other biological assessments (including a histological assessment of the testis from the hybrid males) that demonstrated complete infertility (Table S1).

For instance, in the case of hybrids between lions (*Panthera leo*) and tigers (*P. tigris*), there are numerous anecdotal reports regarding hybrid male sterility in ligers (male lion x female tiger) and the backcross hybrid offspring derived from females (e.g.^40^. Here, we provide histological evidence of sterility based on H&E stained testes from an adult male liger (Fig. S4 panels a and b), and an adult male tiliger (male tiger x female liger; panels c and d). Testes show clear seminiferous tubule degeneration, lined only with Sertoli cells in the liger, and tubule degeneration with germ cell arrest in the tiliger (Fig. S4 panels c, d).

Many additional live hybrid offspring have been reported in the literature than are included in Fig. 1 or in Fig. S1. The relative fertility of the hybrid offspring, however, has not been adequately assessed and we were therefore unable to confidently place them into either category. The framework and threshold values depicted in Fig. 1 allow us to predict the fertility of these offspring given the definitions described above and their placement into Category 1 or 2. These pairs are listed in Table S2 and their relative positions are depicted in Fig. S2.

### Genetic Distance Calculation

Both *CYTB* sequences and full mitogenomes (excluding the control region) of multiple individuals of each species were collected from Genbank (Table S4) and aligned using Clustal Omega version 1.2.4^41^. In order to ensure that none of the sequences was either mislabelled or were nuclear copies of mitochondrial genes, we constructed Neighbour-Joining trees using Geneious version 6.1.8^42^ and removed all individuals that did not fall into monophyletic clades of each species. We first used *pModelTest* version 1.04^43^ to determine the best model for the alignment of each set of sequences for both species. We then calculated pairwise distances between each species pair using RAXML version 8^44^ and FASTTree version 2.1^45^. We also generated raw distance values (i.e. the proportion of sites that differ between each pair of sequences) using the Hamming distance method which sums the number of basepair differences, regardless of whether they are transitions or transversions, and divides that number by the sequence length.

The distances were generated from the *CYTB* (and nuclear gene) alignments for each set of species pairings in fasta file format using a python wrapper to wrap around the programs RAXML, FastTree, pModelTest, and a custom python script written to test Hamming distance of sequences making use of the distance version 0.1.3^46^ and Biopython version 1.68^47^ modules. The script was written for use in version 2.7 of the Terminal application in an OSX environment, and the source code is included at the end of this Supplementary Information. To generate the Hamming distance calculations, the aligned sequences had to be the identical length (meaning all partial sequences were rejected). In addition, gaps in the published sequences were ignored across those loci and across each alignment even if a nucleotide was present at those loci in other sequences to minimise the artificial inflation of the divergence value differences.

### Mann-Whitney test of statistical difference between *CYTB* distance in hybrid categories

The statistical significance of observed differences in mitochondrial genetic distance between groups with reduced hybrid compatibility and those with full compatibility, was tested using the Mann-Whitney U test (p = 5.439e-06) implemented in the R-software package(R Core Team 2013^48^).

## Acknowledgments

We thank Simon Ho, Linda Maxson, Julie Wilson, Andrew Millard, John Hawks, Tom Higham, Kelly Harris, Christian Capelli, Shyam Gopalakirshnan, Montgomery Slatkin, Josh Schraiber, Sam Turvey, Janet Kelso, and Alfred Roca, for advice and discussion. We also thank the Zoological Society and the Bartlet Society for their assistance. GL was supported by the European Research Council (ERC-2013-StG 337574-UNDEAD) and the Natural Environment Research Council (NE/H005269/1 & NE/K005243/1).

## References

1 Currat, M. & Excoffier, L. Modern humans did not admix with Neanderthals during their range expansion into Europe. PLoS Biology 2, e421, doi:10.1371/journal.pbio.0020421 (2004).

2 Serre, D. et al. No evidence of neandertal mtDNA contribution to early modern humans. Early Modern Humans at the Moravian Gate: The MladeČ Caves and their Remains 2, 491–503, doi:10.1007/978-3-211-49294-9_17 (2006).

3 Krings, M. et al. Neandertal DNA Sequences and the Origin of Modern Humans. Cell 90, 19–30, doi:10.1016/S0092-8674(00)80310-4.

4 Green, R. E. et al. A draft sequence of the Neandertal genome. Science (New York, N.Y.) 328, 710–722, doi:10.1126/science.1188021 (2010).

5 Meyer, M. et al. A high-coverage genome sequence from an archaic Denisovan individual. Science 338, 222–226 (2012).

6 Fu, Q. et al. An early modern human from Romania with a recent Neanderthal ancestor. Nature 524, 216–219, doi:10.1038/nature14558 (2015).

7 Keith, A. Ancient Types of Man. (Harper & Brothers, 1911).

8 Sankararaman, S. et al. The genomic landscape of Neanderthal ancestry in present-day humans. Nature 507, 354–357 (2014).

9 Harris, K. & Nielsen, R. The genetic cost of Neanderthal introgression. Genetics 203, 881–891 (2016).

10 Juric, I., Aeschbacher, S. & Coop, G. The Strength of Selection against Neanderthal Introgression. PLOS Genetics 12, e1006340, doi:10.1371/journal.pgen.1006340 (2016).

11 Fu, Q. et al. The genetic history of Ice Age Europe. Nature 534, 200–205 (2016).

12 Cahill, J. A. et al. Genomic evidence of geographically widespread effect of gene flow from polar bears into brown bears. Molecular Ecology 24, 1205–1217, doi:10.1111/mec.13038 (2015).

13 Holliday, T. W. in Neanderthals revisited: New approaches and perspectives 281–297 (Springer, 2006).

14 Holliday, T. W., Gautney, J. R. & Friedl, L. Right for the Wrong Reasons. Current Anthropology 55, 696–724 (2014).

15 JanČúchová-Lásková, J., Landová, E. & Frynta, D. Are genetically distinct lizard species able to hybridize? A review. Current Zoology 61, 155–180 (2015).

16 Edmands, S. Does parental divergence predict reproductive compatibility? Trends in Ecology & Evolution 17, 520–527 (2002).

17 Bradley, R. D. & Baker, R. J. A test of the genetic species concept: cytochrome-b sequences and mammals. Journal of Mammalogy 82, 960–973 (2001).

18 Sánchez-Guillén, R., Córdoba-Aguilar, A., Cordero-Rivera, A. & Wellenreuther, M. Genetic divergence predicts reproductive isolation in damselflies. Journal of evolutionary biology 27, 76–87 (2014).

19 Perelman, P. et al. A Molecular Phylogeny of Living Primates. PLOS Genetics 7, e1001342, doi:10.1371/journal.pgen.1001342 (2011).

20 Presgraves, D. C. & Noor, M. Patterns of postzygotic isolation in Lepidoptera. Evolution 56, 1168–1183 (2002).

21 Price, T. D. & Bouvier, M. M. The evolution of F1 postzygotic incompatibilities in birds. Evolution 56, 2083–2089, doi:10.1111/j.0014-3820.2002.tb00133.x (2002).

22 Coyne, J. A. & Orr, H. A. Patterns of speciation in Drosophila. Evolution, 362–381 (1989).

23 Teeter, K. C. et al. Genome-wide patterns of gene flow across a house mouse hybrid zone. Genome research 18, 67–76 (2008).

24 Song, Y. et al. Adaptive introgression of anticoagulant rodent poison resistance by hybridization between old world mice. Current Biology 21, 1296–1301, doi:10.1016/j.cub.2011.06.043 (2011).

25 Davis, B. W. et al. Mechanisms Underlying Mammalian Hybrid Sterility in Two Feline Interspecies Models. Molecular Biology and Evolution, doi:10.1093/molbev/msv124 (2015).

26 Detlefsen, J. A. Genetic studies on a Cavy species cross. 205 (1914).

27 Thomsen, P. D. et al. Meiotic studies in infertile domestic pig-babirusa hybrids. Cytogenetic and Genome Research 132, 124–128, doi:10.1159/000320421 (2011).

28 Meyer, M. et al. A mitochondrial genome sequence of a hominin from Sima de los Huesos. Nature 505, 403–406 (2014).

29 Meyer, M. et al. A High-Coverage Genome Sequence from an Archaic Denisovan Individual. Science 338, 222–226, doi:10.1126/science.1224344 (2012).

30 Etkind, A. Beyond eugenics: the forgotten scandal of hybridizing humans and apes. Studies in History and Philosophy of Science Part C: Studies in History and Philosophy of Biological and Biomedical Sciences 39, 205–210 (2008).

31 Scally, A. et al. Insights into hominid evolution from the gorilla genome sequence. Nature 483, 169–175 (2012).

32 Hedges, S. B., Marin, J., Suleski, M., Paymer, M. & Kumar, S. Tree of life reveals clock-like speciation and diversification. Molecular biology and evolution, msv037 (2015).

33 Currat, M. & Excoffier, L. Strong reproductive isolation between humans and Neanderthals inferred from observed patterns of introgression. Proceedings of the National Academy of Sciences 108, 15129–15134 (2011).

34 Vernot, B. et al. Excavating Neandertal and Denisovan DNA from the genomes of Melanesian individuals. Science 352, 235–239 (2016).

35 Klein, R. G. Whither the Neanderthals? Science 299, 1525–1527 (2003).

36 Nowell, A. Defining behavioral modernity in the context of Neandertal and anatomically modern human populations. Annual Review of Anthropology 39, 437–452 (2010).

37 Kuhlwilm, M. et al. Ancient gene flow from early modern humans into Eastern Neanderthals. Nature 530, 429–433 (2016).

38 Hill, G. E. Mitonuclear coevolution as the genesis of speciation and the mitochondrial DNA barcode gap. Ecology and Evolution 6, 5831–5842 (2016).

39 Orr, H. A. & Turelli, M. The evolution of postzygotic isolation: accumulating Dobzhansky-Muller incompatibilities. Evolution 55, 1085–1094, doi:10.1111/j.0014-3820.2001.tb00628.x (2001).

40 Gray, A. P. Mammalian hybrids. A check-list with bibliography. [Second edition]. Commonwealth Agricultural Bureaux Slough i-x, 1–262 (1972).

41 Sievers, F. et al. Fast, scalable generation of high-quality protein multiple sequence alignments using Clustal Omega. Molecular Systems Biology 7, 539–539, doi:10.1038/msb.2011.75 (2014).

42 Kearse, M. et al. Geneious Basic: An integrated and extendable desktop software platform for the organization and analysis of sequence data. Bioinformatics 28, 1647–1649, doi:10.1093/bioinformatics/bts199 (2012).

43 Serra, F. pModelTest. (GitHub, 2011).

44 Stamatakis, A. RAxML version 8: a tool for phylogenetic analysis and post-analysis of large phylogenies. Bioinformatics 30, 1312–1313, doi:10.1093/bioinformatics/btu033 (2014).

45 Price, M. N., Dehal, P. S. & Arkin, A. P. FastTree 2--approximately maximum-likelihood trees for large alignments. PloS one 5, e9490, doi:10.1371/journal.pone.0009490 (2010).

46 Meyer, M. distance. (GitHub, 2013).

47 Cock, P. J. a. et al. Biopython: freely available Python tools for compu… [Bioinformatics. 2009] - PubMed result. Bioinformatics (Oxford, England) 25, 1422–1423, doi:10.1093/bioinformatics/btp163 (2009).

48 R: a language and environment for statistical computing GBIF.ORG.

