## Supplementary Materials for "Predicting the viability of archaic human hybrids using a mitochondrial proxy"

### Supplementary Figure Legends

**Figure S1**. This figure is identical to Figure 1 in the main text but includes the numbers associated with each pairwise comparison that are listed in Table S1. The association between the numbers and the pairs (with percent divergence listed in parentheses) are as follows:

1. Papio_hamadryas-Macaca_mulatta (14.2)
2. Sus_scrofa_domesticus-Babyrousa_celebensis (12.9)
3. Peromyscus_truei_comanche-Peromyscus_nasutus (11.8)
4. Panthera_tigris-Panthera_leo (10.1)
5. Papio_hamadryas-Theropithecus_gelada (9.5)
6. Mus_musculus_musculus-Mus_spretus (8.8)
7. Cavia_fulgida-Cavia_porcellus (8.0)
8. Equus_caballus-Equus_asinus (7.7)
9. Pongo_pygmaeus-Pongo_abelii (7.6)
10. Myodes_rutilus-Myodes_glareolus (7.5)
11. Canis_latrans-Canis_aureus (6.5)
12. Canis_latrans-Canis_lupus (6.4)
13. Papio_cynocephalus-Papio_anubis (6.0)
14. Papio_hamadryas-Papio_anubis (5.3)
15. Peromyscus_polionotus-Peromyscus_maniculatus (4.6)
16. Ursus_arctos-Ursus_maritimus (2.4)
17. Mus_musculus_musculus-Mus_musculus_domesticus (2.3)

Hominini hybrid comparisons:

18. Pan_troglodytes-Homo_sapiens_sapiens_modern 11.1
19. Pan_paniscus-Homo_sapiens_sapiens_modern 10.8
20. Homo_sapiens_spp._Denisova-Homo_sapiens_neanderthalensis 2.7
21. Homo_sapiens_spp._Denisova-Homo_sapiens_sapiens_modern 2.5

22. Homo_sapiens_spp._Denisova-Homo_sapiens_sapiens_ancient 2.4
23. Homo_sapiens_neanderthalensis-Homo_sapiens_spp._Sima-de-los-Huesos 2.0
24. Homo_sapiens_spp._Sima-de-los-Huesos-Homo_sapiens_sapiens_modern 1.9
25. Homo_sapiens_spp._Sima-de-los-Huesos-Homo_sapiens_sapiens_ancient 1.8
26. Homo_sapiens_neanderthalensis-Homo_sapiens_sapiens_modern 1.6
27. Homo_sapiens_neanderthalensis-Homo_sapiens_sapiens_ancient 1.6
28. Homo_sapiens_spp._Denisova-Homo_sapiens_spp._Sima-de-los-Huesos 1.3

Felidae hybrid comparisons:
Serval x Cat. Leptailurus_serval-Felis_catus (11.3)
Leopard_Cat x Cat. Prionailurus_bengalensis-Felis_catus (10.9)
Jungle_Cat x Cat. Felis_chaus-Felis_catus (7.5)

**Figure S2**. A comparison of the relative *CYTB* divergence values between those hybrid offspring with known degrees of fertility (green and brown circles, see Figure 1) and those pairs who were able to produce live offspring, but for whom the fertility of their offspring is unknown (white circles). Divergence values are listed on the y-axis as a percentage. Numbers alongside each circle represent specific species pairs and their divergence values are listed in parentheses:

1. Castor_canadensis-Castor_fiber (11.7)
2. Ursus_arctos-Ursus_americanus (11.5)
3. Macaca_nemestrina-Macaca_fascicularis (11.2)
4. Macaca_nemestrina-Macaca_mulatta (10.4)
5. Papio_anubis-Theropithecus_gelada (10.2)
6. Lepus_europaeus-Lepus_timidus (9.8)
7. Macaca_thibetana-Macaca_fascicularis (8.9)
8. Mus_musculus_domesticus-Mus_spretus (8.7)
9. Mustela erminea-Mustela_putorius (8.3)
10. Diceros_bicornis-Ceratotherium_simum_simum (8.2)
11. Macaca_mulatta-Macaca_fascicularis (8.2)
12. Acomys_dimidiatus-Acomys_minous (8.0)

13. Cavia_aperea-Cavia_porcellus (7.9)
14. Loxodonta_africana-Elephas_maximus (7.0)
15. Loxodonta_cyclotis-Loxodonta_africana (4.6)
16. Pan_paniscus-Pan_troglodytes (4.6)
17. Gorilla_beringei_graueri-Gorilla_gorilla_gorilla (4.4)
18. Connochaetes_gnou-Connochaetes_taurinus (2.8)
19. Ceratotherium_simum_cottoni-Ceratotherium_simum_simum (0.9)

The establishment of the framework and the threshold values can be used to predict the relative fertility of the hybrid offspring in cases where there is insufficient experimental information.

**Figure S3**. A comparison of the relative divergence values and pattern between those calculated using CYTB and those using full mitogenomes, and four nuclear genes: ZFY, ZFX, GHR, and CHRNA1. Divergence values are listed on the y-axis for each locus as a percentage. Numbers alongside each circle represent a species pair:

1. Papio_hamadryas-Macaca_mulatta
2. Macaca_nemestrina-Macaca_fascicularis
3. Pan_troglodytes-Homo_sapiens_sapiens
4. Pan_paniscus-Homo_sapiens_sapiens
5. Macaca_nemestrina-Macaca_mulatta
6. Papio_anubis-Theropithecus_gelada
7. Papio_hamadryas-Theropithecus_gelada
8. Macaca_mulatta-Macaca_fascicularis
9. Papio_anubis-Papio_hamadryas
10. Pan_paniscus-Pan_troglodytes

In each case, the distance values of the nuclear genes are smaller relative to those obtained using CYTB as a result of the slower pace of nuclear evolution. Despite this, a clear threshold between the two categories of fertility amongst the hybrid offspring remains.

**Figure S4**. Images of H&E stained testes of an adult male liger (*Panthera leo* x *Panthera tigris*) in panels a and b, and of an adult male tiliger (male tiger x female liger) in panels c and d. The testes show clear seminiferous tubule degeneration, lined only with Sertoli cells in the liger, and tubule degeneration with germ cell arrest in the tiliger.

**Source code for a custom Python version 2.7 terminal program to calculate pair-wise Hamming distances between the sequences contained within a Fasta file.**

```
#!/usr/bin/python

#import all python 2.7 native modules
import operator, StringIO, itertools, sys, math, os

#import all non-native modules
import distance, Bio

#import various classes from modules
from Bio import SeqIO
from Bio import AlignIO
from Bio.Align import AlignInfo
from itertools import izip, imap
from os import path

#disable screen blanking and the terminal cursor
os.system('setterm -cursor off')

#calculate hamming distance between two strings
def hamming(str1, str2):
    assert len(str1) == len(str2)
    ne = str.__ne__
    ne = operator.ne
    return sum(imap(ne, str1, str2))

#count gaps in the consensus sequence
def count_gaps(consensus):
    b = 0
    for a in range(0,len(consensus)):
        if consensus[a] == '-':
            b +=1
```

```
    return b

#build a consensus sequence from two sequences
def consensus_seq(str1,str2):
    consensus_string = []
    for a, b in zip(range(0,len(str1)),range(0,len(str2))):
        if str1[a] != str2[b]:
            if str1[a]== '-' or str2[b] == '-':
                consensus_string.append('-')
            else:
                consensus_string.append('N')
        else:
            consensus_string.append(str1[a])
    return ''.join(consensus_string)

#compare the gaps locations between pairwise sequences
def compare_gaps(str1,str2):
    gaps1 = 0
    gaps2 = 0
    gaps3 = 0
    for bp1, bp2 in zip(str1,str2):
        if bp1 == "-" and bp1 == bp2:
            gaps1 +=1
        if bp1 == "-" and bp1 != bp2:
            gaps2 +=1
        if bp2 == "-" and bp2 != bp1:
            gaps3 +=1
    return [gaps1,gaps2,gaps3]

#clearout the command line so the progress counter remains in the same
location on the screen
def restart_line():
        sys.stdout.write('\r')
        sys.stdout.flush()

#grab user defined input file from terminal
fasta_file = sys.argv[1]

#create a file to output raw distances to
raw_distance_file = open(sys.argv[2],'w')

```

```
#parse the fasta file using biopython to create iterable fasta object
sequences = SeqIO.parse(open(fasta_file),"fasta")

#define a list to append sequence data to
sequence_list = []

#iterate over fasta sequence objects and add their id and sequence to a
line separated by a comma
for record in sequences:
    sequence_list.append(record.id + ',' + record.seq)

#create an array containing all the possible pairwise combinations of all
the sequences in the sequence list
pairwise_sequences = itertools.combinations(sequence_list,2)

#print a spacing line in the terminal output
print""

#count the number of pairwise comparisons and put into a variable
for i, item in enumerate(itertools.combinations(sequence_list,2)):
no_pairwise_seqs = i

#for each pair_wise comparison between sequences
for i, item in enumerate(pairwise_sequences):

    #create an output file like class that can be written to and read from
    output = StringIO.StringIO()

    #format the two strings for comparison into fasta format in two string
variables
    str1 = ">"+item[0].split(',')[0]+"\n"+item[0].split(',')[1]+"\n"
    str2 = ">"+item[1].split(',')[0]+"\n"+item[1].split(',')[1]+"\n"

    #create a temporary fasta file on the hard drive
    temp_fasta = open("/home/richard/Documents/temp_fasta.fasta","w")

    #write the fasta formatted sequences to the output class
    output.write(str1+'\n')
    output.write(str2+'\n')

    #store the value of the output class to a variable
```

```
    contents = output.getvalue()

    #close the output class removing it from memory
    output.close()

    #write the contents of the variable to the temporary fasta file
    temp_fasta.write(contents)

    #close the fasta file removing it from memory and saving the changes
    temp_fasta.close()

    #grab temp file name and put into a variable
    temp_fasta = "/home/richard/Documents/temp_fasta.fasta"

    #create an alignment using the sequences in the temp file
    alignment = AlignIO.read(open(temp_fasta),"fasta")

    #summary_align = AlignInfo.SummaryInfo(alignment)

    #grab the consensus sequence from the pairwise aligment
    consensus = consensus_seq(alignment[0].seq,alignment[1].seq)

    #count the gaps in each sequence including the consensus
    gaps3 = float(count_gaps(alignment[1].seq))
    gaps2 = float(count_gaps(alignment[0].seq))
    gaps1 = float(count_gaps(consensus))

    #use the hammings distance method from the distance module in both
directions for both sequences
    dist1 = distance.hamming(alignment[0].seq,alignment[1].seq)

    #compare the locations of gaps between the pairwise seuqneces and their
consensus sequence to calculate total genuine gaps
    total_gaps = compare_gaps(alignment[0].seq,alignment[1].seq)

    #calculate the properdistance between sequences excluding gaps
    answer = float(dist1)-(total_gaps[1]+total_gaps[2])

    #convert the number of distances into a floating point number that can
be converted to a percentage difference later
```

```
    percent_count = "%.2f" % round(i/float(no_pairwise_seqs)*100,2)

    #write an update percentage to the command line to indicate progress
through the different combinations
    sys.stdout.write( str(percent_count) + "% complete...")
    sys.stdout.flush()
    restart_line()

    #write the pairwise distances to an output file to be used later
    raw_distance_file.write(alignment[0].id + "  " + alignment[1].id + "  " +
str(answer/len(consensus))+"\n")

#print a spacing line in the terminal output
print ""

#close the file containing the pairwise distances
raw_distance_file.close()

#restore the default screen blanking and terminal cursor settings
os.system('setterm -cursor on')
```

**Supplementary Table 1**. A list of species pairs known to produce viable offspring with the exception of the first pair (labelled Hybrid Pair 0) that was unable to produce offspring at all. Green and Brown shading represent pairs listed in Category 1 and Category 2 respectively. Pairs of species with no background colour correspond to the inter-hominini comparisons discussed in the text. All pairs are depicted on Figure 1 and are listed by number in Supplementary Figure 1. The percent distances are those calculated using raw Hammings distances calculations of *CYTB* divergence described in the text and Supplementary Information. 95% confidence intervals were calculated by averaging pairwise distances between species groups and reported as standard error.

| Pair | Species (Binomial) | Species (Common) | KT (2n) | % Dist | St.Err | # Seqs | Seq Len | Hybrid Notes |
|---|---|---|---|---|---|---|---|---|
| 0 | *Oryctolagus cuniculus* | European Rabbit | 44(1) | 17.4 | 4.50E-04 | 4 | 1140 | Breeding experiments yielded no progeny. Attempts at crossing hares and rabbits reciprocally by natural means were not successful. The hare would not consent to mate with a rabbit. Sperm in which lively movement could be observed under the microscope was taken from the vasa deferentia of hares, diluted with dextrose solution, and then injected into the uterus of a female rabbit which had just copulated with a sterile male rabbit. In 38 cases, the result was entirely negative. The authors conclude that it is impossible to get hares to mate naturally with rabbits, and even if this were to occur under exceptional circumstances, no hybrid offspring would result because of the evident inability of the rabbit egg to be fertilized by hare sperm(2). |
| | *Lepus timidus* | Mountain Hare | 48(3) | | | 20 | | |

| Pair | Species (Binomial) | Species (Common) | KT (2n) | % Dist | St.Err | # Seqs | Seq Len | Hybrid Notes |
|---|---|---|---|---|---|---|---|---|
| 1 | *Macaca mulatta* | Rhesus Macaque | 42(4) | 14.17 | 1.25E-04 | 108 | 1141 | In Leningrad, this cross was an intergeneric hybrid between a baboon and rhesus monkey(5)). In the late 1970s, so-called Rheboons were bred from female baboons and male macaques at the Southwest Foundation for Biomedical Research in San Antonio, Texas. Four infants (two male and two female) survived the newborn period. A long-term male Rheboon survivor (18 years) underwent genetic analysis and was shown to have a karyotype of 2n=42. Analysis of the male hybrid's semen indicated he was completely infertile with an absence of spermatazoa(4). |
| | *Papio hamadryas* | Hamadryas Baboon | 42(4) | | | 2 | | |
| 2 | *Sus domesticus* | Domestic Pig | 38(6) | 12.89 | 4.43E-04 | 4 | 1140 | At Copenhagen Zoo in 2006, a babirusa boar and a domestic sow produced five piglets of both sexes. Of three surviving piglets, a male hybrid had a karyotype of 2n = 38. Analysis of semen revealed meotic arrest after prophase I. Some female oocytes appeared normal(6). |
| | *Babyrousa celebensis* | Babirusa | 38(6) | | | 67 | | |
| 3 | *Peromyscus truei comanche* | Palo Duro Mouse | 48(7) | 11.85 | 9.62E-04 | 67 | 1144 | Gray(8) reported that male F1 hybrids are sterile. Six F1 males were backcrossed to each parent species. When the cross was with *P.truei*, one out of six matings was successful which resulted in one male and one female offspring. When the cross was with *P.nastus*, no offspring resulted. Several male hybrids produced either no or deformed spermatozoa in their semen(9),(10). |
| | *Peromyscus nasutus* | Northern Rock Mouse | 48(7) | | | | | |

| Pair | Species (Binomial) | Species (Common) | KT (2n) | % Dist | St.Err | # Seqs | Seq Len | Hybrid Notes |
|---|---|---|---|---|---|---|---|---|
| 4 | *Panthera tigris* | Tiger | 38(11) | 10.07 | 5.90E-04 | 4 | 1137 | A male lion and a female F1 hybrid Liger produced a viable female offspring(8)). In 2012, in the Novosibirsk Zoo in Russia, a liger F1 female and a male African lion produced an offspring (a Liliger) named Kiara. The same pairing has produced three more female hybrids(12). Reciprocal male F1 hybrids have been shown to be sterile, with seminiferous tubule degeneration in ligers, and germ cell arrest in tiliger males (Supplementary Text). |
| | *Panthera leo* | Lion | 38(13) | | | 4 | | |
| 5 | *Papio hamadryas* | Hamadryas Baboon | 42(14) | 9.49 | 2.29E-04 | 40 | 1141 | F1 hybrids (male and female) viable hybrids have been produced. Though an F1 male was described as “fertile”, pregnancies with F1 hybrid females did not result in viable offspring(15). |
| | *Theropithecus gelada* | Gelada | 42(15) | | | 10 | | |

| Pair | Species (Binomial) | Species (Common) | KT (2n) | % Dist | St.Err | # Seqs | Seq Len | Hybrid Notes |
|---|---|---|---|---|---|---|---|---|
| 6 | *Mus musculus musculus* | House Mouse | 40(16) | 8.85 | 1.09E-04 | 4 | 1141 | These two species are partially sympatric in Africa and Europe. Male and some female hybrids are infertile, depending on the direction of the cross. Experimental introgression has transferred an anticoagulant rodent poison resistance gene from *M.spretus* into *M.domesticus* and this has taken place naturally in wild populations in Germany and Spain(17). F1 males are infertile and F1 females can be backcrossed to male *M.spretus* males, but have reduced reproductive capacity. Five of 16 male backcross hybrids were fertile and the rest were infertile. Fertility was confirmed by breeding them with backcrossed females(18). Evidence of 6.5% introgressed *M.spretus* genomic elements were found in the C57BL/BJ strain used in breeding experiments, indicating historical gene flow between the two species(19). This strain also has a Y-chromosone haplogroup consistent with Asian *Mus musculus musculus* and not *Mus musculus domesticus*(20). |
| | *Mus spretus* | Algerian Mouse | 40(16) | | | 4 | | |
| 7 | *Cavia fulgida* | Shiny guinea pig | 64(21) | 8.02 | 3.38E-04 | 2 | 1140 | Breeding experiments produced sterile F1 males. However, the 3rd generation of backcrosses to *C. porcellus* generated a hybrid with restored fertility(22). |
| | *Cavia porcellus* | Guinea pig | 64(21) | | | 2 | | |

| Pair | Species (Binomial) | Species (Common) | KT (2n) | % Dist | St.Err | # Seqs | Seq Len | Hybrid Notes |
|---|---|---|---|---|---|---|---|---|
| 8 | *Equus caballus* | Horse | 64(23) | 7.73 | 5.49E-05 | 9 | 1140 | In China in 1981, a fertile female mule gave birth to a foal sired by a male donkey, which was confirmed by genetic testing. The foal displayed a unique karyotype not previously described in a hybrid. The mother had regular oestrous cycles and the resultant foal had a 2n karyotype of 62, the same as a donkey and showed strong oestrus behaviour. Analysis showed that enough of the oocytes survive meiosis to allow oestrus to occur at regular intervals(24). |
| | *Equus asinus* | Donkey | 62(23) | | | 9 | | |
| 9 | *Myodes rutilus* | Northern red-backed vole | 56(25) | 7.54 | 1.45E-05 | 9 | 1140 | This pair can produce hybrids when crossed under laboratory conditions(26). Though many published papers report the finding that bank voles possess red-vole mitochondrial DNA (mtDNA) in different sites of the sympatric range (27), hybrid males are sterile, whereas hybrid females are fertile. |
| | *Myodes glareolus* | Bank Vole | 56(25) | | | 9 | | |
| 10 | *Pongo pygmaeus* | Bornean Orangutan | 48(28) | 7.60 | 2.92E-04 | 3 | 1140 | F1 hybrids have been bred for up to four generations amongst themselves without any indication of reduced fertility(29), though a high level of failure of F1 births has been reported(8). As of 1990, 29% of the total US captive population of 300 individuals was described as F1 hybrids(30). |
| | *Pongo abelii* | Sumatran Orangutan | 48(28) | | | 3 | | |

| Pair | Species (Binomial) | Species (Common) | KT (2n) | % Dist | St.Err | # Seqs | Seq Len | Hybrid Notes |
|---|---|---|---|---|---|---|---|---|
| 11 | *Canis latrans* | Coyote | 78(31) | 6.49 | 1.52E-03 | 5 | 1140 | The Nuremburg Zoo (Germany) reported a male and a female F2 hybrid resulting from the matings of F1 hybrid female and a male coyote. Additional *C.aureus* x *C.latrans* reciprocal crosses have been reported, and some hybrids have been demonstrated to be fertile(8). |
| | *Canis aureus* | Golden Jackal | 78(32) | | | 5 | | |
| 12 | *Canis latrans* | Coyote | 78(31) | 6.38 | 3.31E-04 | 5 | 1140 | Coyote DNA was first reported in *C.lupus* populations in the Great Lakes in the 1990s. There is evidence of introgression with Eastern and Western wolves and domestic dogs with coyotes. Male-mediated introgression is thought to be the most common, with Eastern male wolves coupling with coyote females. Every 1 in 70 coyotes have wolf mitochondrial genomes. Rather than sterility, this sex bias is thought to be size mediated with males of the larger species mating with females of the smaller species(33). During controlled experiments, a male wolf and female coyote produced two hybrid litters of five pups each(8). In 2014, it was reported that one western coyote female successfully produced six viable hybrid offspring with a male western grey wolf through artificial insemination(34). In 2017, these F1 hybrids were reported to have survived the first four years and subsequently have produced F2 hybrids by mating with each other. (35). |
| | *Canis lupus* | Wolf | 78(31) | | | 5 | | |

| Pair | Species (Binomial) | Species (Common) | KT (2n) | % Dist | St.Err | # Seqs | Seq Len | Hybrid Notes |
|---|---|---|---|---|---|---|---|---|
| 13 | *Papio anubis* | Olive Baboon | 42(36) | 5.96 | 2.51E-04 | 40 | 1141 | Multiple natural hybrid zones exist within Kenya and Eastern Africa in general. Both species share unusually similar mtDNA haplotypes. There is only a single nucleotide substitution difference implying introgression of the mtDNA from one species to the other and at least partial fertility(37). In contact zones, baboon taxa hybridize readily, producing viable and fertile hybrid offspring(38). All baboon allotaxa appear to be capable of producing viable and fertile offspring when crossed(37, 39-45). A hybrid was also produced In Chessington Zoo(46). Captive F1 and B1 hybrids have also been produced at the Southwest Foundation for Biomedical Research (SFBR)(47). F1 hybrids have also been reported to be capable of producing F2 hybrids(48). |
| | *Papio cynocephalus* | Yellow Baboon | 42(14) | | | 40 | | |
| 14 | *Papio anubis* | Olive Baboon | 42(36) | 5.30 | 9.68E-05 | 40 | 1141 | A natural hybrid zone exists in Ethiopia's Awash National Park. A group of hybrid individuals was studied in detail and found to be a blend of elements of both species(49). Like the Olive and Yellow baboons, all baboon allotaxa appear to be capable of producing viable and fertile offspring when crossed(37, 39-45). Fully fertile hybrids are more likely when the mother is *P.hamadryas*(50). |
| | *Papio hamadryas* | Hamadryas Baboon | 42(14) | | | 40 | | |

| Pair | Species (Binomial) | Species (Common) | KT (2n) | % Dist | St.Err | # Seqs | Seq Len | Hybrid Notes |
|---|---|---|---|---|---|---|---|---|
| 15 | *Peromyscus polionotus* | Oldfield Mouse | 48(51) | 4.65 | 9.53E-05 | 67 | 1141 | These species will hybridise in the lab(52), though the F1 hybrids are more likely to be fully fertile when the female deer mouse is the mother(52, 53). The F1 hybrids are almost as fertile in both directions (back crossing or with each other), though there is a slight reduced fertility rate in crosses(54). |
| | *Peromyscus maniculatus* | Deer Mouse | 48(51) | | | 67 | | |
| 16 | *Ursus arctos* | Brown Bear | 74(28) | 2.42 | 3.53E-05 | 6 | 1140 | In 1876 in a zoo in Stuttgart, Germany, a female European brown bear was mated with a male polar bear resulting in a hybrid cub. A further three births with this pair were reported. The young were subsequently bred with each other to produce cubs(55). The bears of ABC Island have evidence of introgression with polar bears, because they have polar bear mtDNA and 6.5% introgression of X-chromosone DNA from polar bears(56). There appears to be evidence of an 8.8% genetic contribution in populations of brown bear from polar bears, however, currently there is no evidence of the reverse, which is likely the result of ecological selection rather than a barrier to gene flow(57). |
| | *Ursus maritimus* | Polar Bear | 74(28) | | | 6 | | |

| Pair | Species (Binomial) | Species (Common) | KT (2n) | % Dist | St.Err | # Seqs | Seq Len | Hybrid Notes |
|---|---|---|---|---|---|---|---|---|
| 17 | *Mus musculus musculus* | House Mouse | 40(16) | 2.30 | 6.44E-05 | 4 | 1141 | A hybrid zone exists between Eastern and Western Europe from Denmark to Bulgaria(58). Hybrids have been bred in lab settings that are fully fertile. Occasional male and female sterility can result due to polymorphisms in genes within the X-chromosome causing asymmetric autosomal pairing during meiosis(59). |
| | *Mus musculus domesticus* | Domestic House Mouse | 40(16) | | | 4 | | |
| 18 | *Homo sapiens* | Modern Human | 46(28) | 11.07 | 1.05E-04 | 34 | 1140 | Experiments were carried out by the Soviet Scientist Ilya Ivanovich Ivanov during the 1920s, in which female chimps were injected with human sperm. None of the inseminations resulted in pregnancy. He later decided to try and impregnate female human volunteers with ape sperm (chimp and orangutan) but the captured apes failed to flourish except for the orangutan. Before Ivanov could carry out the experiment with the remaining ape, the ape suffered from a brain haemorrhage (60, 61). |
| | *Pan troglodytes* | Chimpanzee | 48(28) | | | 40 | | |
| 19 | *Homo sapiens* | Modern Human | 46(28) | 10.83 | 5.57E-05 | 34 | 1140 | It is uncertain if this experiment has been attempted. |
| | *Pan paniscus* | Bonobo | 48(28) | | | 39 | | |
| 20 | *Homo sapiens spp.* Denisova | Denisovan | 46(62) | 2.68 | 1.89E-04 | 4 | 1140 | A complete nuclear genome from a Neanderthal woman found in Siberia was sequenced and analysed for introgression. When compared to published sequences of Denisovan DNA, it was found that 0.5% of the Neanderthal genome was introgressed into the published Denisovan genomes indicating historic hybridisation of the two species(63). |
| | *Homo sapiens neanderthalensis* | Neanderthal | 46(62) | | | 20 | | |

| Pair | Species (Binomial) | Species (Common) | KT (2n) | % Dist | St.Err | # Seqs | Seq Len | Hybrid Notes |
|---|---|---|---|---|---|---|---|---|
| 21 | *Homo sapiens spp.* Denisova | Denisovan | 46(62) | 2.50 | 1.80E-04 | 4 | 1140 | A distal manual phalanx recovered from Denisova Cave in the Altai mountains produced a 1.9x genome. Analysis showed introgression of Denisovan DNA into modern day Melansesians and Austronesians(64). |
| | *Homo sapiens sapiens* | Modern human | 46(28) | | | 34 | | |
| 22 | *Homo sapiens spp.* Denisova | Denisovan | 46(62) | 2.40 | 3.43E-04 | 4 | 1140 | See pair 21. |
| | *Homo sapiens sapiens* | Ancient Modern human | 46(28) | | | 8 | | |
| 23 | *Homo sapiens neanderthalensis* | Neanderthal | 46(62) | 2.05 | 1.73E-04 | 20 | 1140 | Mitochondrial sequences have been obtained for an individual currently classified as *H. heidelburgensis*(65). Nuclear genes later sequenced from the same sample indicates that this individual is a basal to Neanderthals(66). There is currently no evidence of introgression into other hominins. |
| | *Homo sapiens .spp* Sima de los Huesos | Proto Neanderthal | 46(62) | | | 2 | | |
| 24 | *Homo sapiens .spp* Sima de los Huesos | Proto Neanderthal | 46(62) | 1.88 | 1.53E-04 | 2 | 1140 | See pair 23. |
| | *Homo sapiens sapiens* | Modern human | 46(28) | | | 34 | | |
| 25 | *Homo sapiens .spp* Sima de los Huesos | Proto Neanderthal | 46(62) | 1.79 | 2.51E-04 | 2 | 1140 | See pair 23. |
| | *Homo sapiens sapiens* | Ancient Modern human | 46(28) | | | 8 | | |
| 26 | *Homo sapiens neanderthalensis* | Neanderthal | 46(62) | 1.61 | 4.28E-05 | 20 | 1140 | Evidence of introgression of up to 9% of the X-chromosome from Neanderthals into non-African populations of modern humans (67). There is also an average of 1.38% and 1.15% of whole genome introgression from Neanderthals into East Asian and European populations respectively(68). |
| | *Homo sapiens sapiens* | Modern human | 46(28) | | | 34 | | |
| 27 | *Homo sapiens neanderthalensis* | Neanderthal | 46(62) | 1.55 | 6.07E-05 | 20 | 1140 | See pair 26. |
| | *Homo sapiens sapiens* | Ancient Modern human | 46(28) | | | 8 | | |
| 28 | *Homo sapiens spp.* Denisova | Denisovan | 46(62) | 1.29 | 5.43E-04 | 4 | 1140 | See pair 23. |
| | *Homo sapiens .spp* Sima de los Huesos | Proto Neanderthal | 46(62) | | | 2 | | |

1. Schröder J & van der Loo W (1979) Comparison of karyotypes in three species of rabbit: Oryctolagus cuniculus, Sylvilagus nuttallii and S. idahoensis. *Hereditas* 91:27-30.
2. Castle WE (1925) The Hare-Rabbit, A Study in Evolution by Hybridization. *The American Naturalist* 59:280-283.
3. Chapman Ja & Flux JEC (1990) Rabbits, hares and pikas: status survey and conservation action plan.61-94.
4. Moore CM*, et al.* (1999) Cytogenetic and fertility studies of a rheboon, rhesus macaque (Macaca mulatta) x baboon (Papio hamadryas) cross: Further support for a single karyotype nomenclature. *American Journal of Physical Anthropology* 110:119-127.
5. Voronin LG (1950) In To Aftica for Mokys.
6. Thomsen PD*, et al.* (2011) Meiotic studies in infertile domestic pig-babirusa hybrids. *Cytogenetic and Genome Research* 132:124-128.
7. Hsu TC & Arrighi FE (1966) Chromosomal evolution in the genus Peromyscus (Cricetidae, Rodentia). *Cytogenetics* 5:355-359.
8. Gray AP (1972) Mammalian hybrids. A check-list with bibliography. [Second edition]. *Commonwealth Agricultural Bureaux Slough i-x*:1-262.
9. Blair WF (1943) Populations of the deer-mouse and associated small mammals in the mesquite association of southern New Mexico. *Contributions of the Laboratory of Vertebrate Biology, University of Michigan* 21:1-40.
10. Tamsitt JR (1958) The Baculum of the Peromyscus truei Species Group. *Journal of Mammalogy* 39:598.
11. Vinogradov AE (1998) Genome size and GC-percent in vertebrates as determined by flow cytometry: The triangular relationship. *Cytometry* 31:100-109.
12. Andreassi K (2012) ""Liliger" Born in Russia No Boon for Big Cats". *National Geographic.*
13. Tian Y, Nie W, Wang J, Ferguson-Smith Ma, & Yang F (2004) Chromosome evolution in bears: Reconstructing phylogenetic relationships by cross-species chromosome painting. *Chromosome Research* 12:55-63.
14. Chiarelli B, Koen AL, & Ardito G (1979) Comparative karyology of primates.
15. Markarjan DS, Isakov EP, & Kondakov GI (1974) Intergeneric hybrids of the lower (42-chromosome) monkey species of the Sukhumi monkey colony. *Journal of Human Evolution* 3:247-255.
16. Silver LM (1995) Mouse Genetics. Concepts and Applications.376.
17. Song Y*, et al.* (2011) Adaptive introgression of anticoagulant rodent poison resistance by hybridization between old world mice. *Current Biology* 21:1296-1301.
18. Hale DW, Washburn LL, & Eicher EM (1993) Meiotic abnormalities in hybrid mice of the C57BL/6J x Mus spretus cross suggest a cytogenetic basis for Haldane's rule of hybrid sterility. *Cytogenetics and cell genetics* 63:221-234.
19. Rikke BA, Zhao Y, Daggett LP, Reyes R, & Hardies SC (1995) Mus spretus LINE-1 sequences detected in the Mus musculus inbred strain C57BL/6J using LINE-1 DNA probes. *Genetics* 139:901-906.
20. Tucker PK, Lee BK, Lundrigan BL, & Eicher EM (1992) Geographic origin of the Y Chromosomes in ?old? inbred strains of mice. *Mammalian Genome* 3:254-261.
21. Gava A, dos Santos MB, & Quintela FM (2011) A new karyotype for Cavia magna (Rodentia: Caviidae) from an estuarine island and C. aperea from adjacent mainland. *Acta Theriologica* 57:9-14.
22. Detlefsen JA (1914) Genetic studies on a *Cavy* species cross. 205.
23. Benirschke K, Brownhill LE, & Beath MM (1962) Somatic chromosomes of the horse, the donkey and their hybrids, the mule and the hinny. *Journal of reproduction and fertility* 4:319-326.
24. Rong RH, Cai HD, Yang XQ, & Wei J (1985) Fertile mule in China and her unusual foal. *Journal of the Royal Society of Medicine* 78:821-825.
25. KANEKO Y, NAKATA K, SAITOH T, STENSETH NC, & BJØRNSTAD ON (1998) The Biology of the Vole Clethrionomys rufocanus: a Review. *Researches on population ecology* 40:21-37.
26. Osipova OV & Soktin AA (2006) Bank and red vole hybridization under experimental conditions. *Doklady Biological Sciences* 410:381-383.
27. Potapov SG*, et al.* (2007) Transfer of mitochondrial genome of the northern redbacked vole (Clethrionomys rutilus) to the bank vole (C. glareolus) in northwestern Europe. *Doklady biological sciences : proceedings of the Academy of Sciences of the USSR, Biological sciences sections / translated from Russian* 417:435-438.
28. Kaplan AR (1971) Atlas of Mammalian Chromosomes - Hsu,Tc and Benirschke,K. *Social Biology* 18:325.

29. Zhi L, *et al.* (1996) Genomic differentiation among natural populations of orang-utan (Pongo pygmaeus). *Current biology : CB* 6:1326-1336.
30. Perkins & Maple TL (1990) North American orangutan species survival plan. *Zoo Biology* 9:135.
31. Snyder DA & Hungerford LK (1966) Chromosomes of a European wolf (Canis lupus) and of Bactrian Camel (Camelus bactrianus). *Mammal Chromosomes Newsletter* 20.
32. Datta M (1979) Evolutionary Significance of the Chromosome Numbers in Mammals: a point of view. in *Comparative Karyology of Primates* (World Anthropology - De Gruyter), p 75.
33. Monzõn J, Kays R, & Dykhuizen DE (2014) Assessment of coyote-wolf-dog admixture using ancestry-informative diagnostic SNPs. *Molecular Ecology* 23:182-197.
34. Mech LD, *et al.* (2014) Production of Hybrids between Western Gray Wolves and Western Coyotes. *PLoS ONE* 9:e88861.
35. Mech LD, *et al.* (2017) Studies of wolf x coyote hybridization via artificial insemination. *PLOS ONE*.
36. Dutrillaux B, Biemont MC, Viegas-Pequignot E, & Laurent C (1979) Comparison of the karyotypes of four Cercopithecoidae: Papio papio, P. anubis, Macaca mulatta, and M. fascicularis. *Cytogenetics and cell genetics* 23:77-83.
37. Newman TK JC, Rogers J. (2004) Mitochondrial phylogeny and systematics of baboons (Papio). Am J Phys Anthropol 124:17–27. *American Journal of Physical Anthropology* 124:17-27.
38. Charpentier MJE, *et al.* (2012) Genetic structure in a dynamic baboon hybrid zone corroborates behavioural observations in a hybrid population. *Molecular Ecology* 21:715-731.
39. Jolly CJ (1993) Species, subspecies and baboon systematics. in *Species, species concepts and primate evolution* (Wiley, New York), pp 67-107.
40. Maples WR & McKern TW (1967) A preliminary report on classification of the Kenya baboon. in *Vartborg, H. (ed.),The Baboon in Medical Research vol.2* (University of Texas Press, Austin), pp 13-22.
41. Nagel U (1973) A comparison of anubis baboons, hamadryas baboons and their hybrids at a species border in Ethiopia. *Folia primatologica; international journal of primatology* 19:104-165.
42. Phillips-Conroy JE & Jolly CJ (1986) Changes in the structure of the baboon hybrid zone in the Awash National Park, Ethiopia. *American Journal of Physical Anthropology* 71:337-350.
43. Alberts SC & Altmann J (2001) Immigration and Hybridization Patterns of Yellow and Anubis Baboons In and Around Amboseli, Kenya. *American Journal of Primatology J. Primatol* 53:139-154.
44. Jolly CJ & Phillips-Conroy JE (2007) Ecology, history and society as determinants of hybrid zone structure in baboons. *American Journal of Physical Anthropology* 132:138.
45. Tung J, Charpentier MJE, Garfield DA, Altmann J, & Alberts SC (2008) Genetic evidence reveals temporal change in hybridization patterns in a wild baboon population. *Molecular ecology* 17:1998-2011.
46. Chessington-Zoo (1970) Chessington Zoo Official Guide.24.
47. Ackermann RR, Rogers J, & Cheverud JM (2006) Identifying the morphological signatures of hybridization in primate and human evolution. *Journal of human evolution* 51:632-645.
48. Ackermann RR, Schroeder L, Rogers J, & Cheverud JM (2014) Further evidence for phenotypic signatures of hybridization in descendant baboon populations. *Journal of Human Evolution* 76:54-62.
49. Bergman TJ & Beehner JC (2004) Social System of a Hybrid Baboon Group (<i>Papio anubis Ã— P. hamadryas</i>). *International Journal of Primatology* 25 %W http:1313-1330.
50. Chiarelli aB & Capanna E (1973) Cytotaxonomy and vertebrate evolution.xv, 783.
51. Dixon LK, Nelson BA, & Priest RL (1984) Chromosome differences in Peromyscus maniculatus populations at different altitudes in Colorado. *Genetica* 52-53:63-68.
52. Watson ML (1942) Hybridization Experiments between Peromyscus Polionotus and Peromyscus Maniculatus. *Journal of Mammalogy* 23:315.
53. Joyner CP, Myrick LC, Crossland JP, & Dawson WD (1998) Deer Mice As Laboratory Animals. *ILAR Journal* 39:322-330.
54. Dawson WD (1965) Fertility and Size Inheritance in a Peromyscus Species Cross. *Evolution* 19:44.
55. Scherren H (1907) 4. Some Notes on Hybrid Bears. *Proceedings of the Zoological Society of London* 77:431-435.
56. Cahill Ja, *et al.* (2013) Genomic Evidence for Island Population Conversion Resolves Conflicting Theories of Polar Bear Evolution. *PLoS Genetics* 9:1-8.

57. Cahill JA*, et al.* (2015) Genomic evidence of geographically widespread effect of gene flow from polar bears into brown bears. *Molecular Ecology* 24(6):1205-1217.
58. Alibert P, Renaud S, Dod B, Bonhomme F, & Auffray JC (1994) Fluctuating asymmetry in the Mus musculus hybrid zone: a heterotic effect in disrupted co-adapted genomes. *Proceedings. Biological sciences / The Royal Society* 258:53-59.
59. Bhattacharyya T*, et al.* (2014) X Chromosome Control of Meiotic Chromosome Synapsis in Mouse Inter-Subspecific Hybrids. *PLoS Genetics* 10:e1004088.
60. Rossiianov K (2002) Beyond Species: Il'ya Ivanov and His Experiments on Cross-Breeding Humans with Anthropoid Apes. *Science in Context* 15:277-316.
61. Pain S (2008) Blasts from the past: The Soviet ape-man scandal. in *New Scientist*.
62. Meyer M*, et al.* (2012) A High-Coverage Genome Sequence from an Archaic Denisovan Individual. *Science* 338:222-226.
63. Prüfer K*, et al.* (2014) The complete genome sequence of a Neanderthal from the Altai Mountains. *Nature* 505:43-49.
64. Reich D*, et al.* (2010) Genetic history of an archaic hominin group from Denisova Cave in Siberia. *Nature* 468:1053-1060.
65. Meyer M*, et al.* (2014) A mitochondrial genome sequence of a hominin from Sima de los Huesos. *Nature* 505(7483):403-406.
66. Meyer M*, et al.* (2016) Nuclear DNA sequences from the Middle Pleistocene Sima de los Huesos hominins. *Nature* 531:504-507.
67. Yotova V*, et al.* (2011) An X-Linked Haplotype of Neandertal Origin Is Present Among All Non-African Populations. *Mol Biol Evol* 28:1957-1962.
68. Sankararaman S*, et al.* (2014) The genomic landscape of Neanderthal ancestry in present-day humans. *Nature* 507(7492):354-357.

Table 2. A list of hybrids known to produce offspring where there is little to no data on subsequent hybrid fertility

| Pair | Species (Binomial) | Species (Common) | KT (2n) | % Dist | St.Err | # Seqs | Seq Len | PC | Hybrid Notes |
|---|---|---|---|---|---|---|---|---|---|
| 1 | *Castor canadensis* | American Beaver | 40(1) | 11.68 | 3.09E-04 | 1 | 1140 | 2 | Russian breeding experiments produced a still-born hybrid between a female Eurasian and male Canadian beaver that possessed intermediate features. 27 breeding experiments prior to this had resulted in matings but no pregnancies. The still birth of the experimental hybrid could have been due to the mother being a first-time parent (which is not uncommon with beavers)(2). Three populations of Eurasian beavers were genetically analysed and the Austrian population had three individuals that appeared to be intermediate between the two species of beaver. Hybridisation was initially dismissed due to the work of Lavrov and the assumption that karyotype acts as a barrier to gene flow, which may not be the case(3). |
| | *Castor fiber* | Eurasian Beaver | 48(1) | | | 11 | | | |
| 2 | *Ursus arctos* | Brown Bear | 74(4) | 11.49 | 5.45E-05 | 34 | 1140 | 2 | *Ursus americanus* x *Ursus arctos* crosses have likely occurred in both directions. Three hybrids of both sexes born in the London Zoological Gardens in 1859, all died(5). A *U. arctos* x *U. americanus* female produced offspring at St. Paul, USA in 1969 when bred to male grizzly bear *U. arctos horribilis*. A *U. americanus* x *U. a. lasiotus*, probably male *U.a. losiotusl* x female *U .americanus*, produced hybrid offspring in Pittsburgh in 1963, 1965 and 1967(6). In 2012, a suspect male hybrid bear called "Ben" (*U. americanus* x *U. arctos*) was rescued from a roadside Zoo in North Carolina and transferred by Fedex via "Bearforce One" to a sanctuary in Northern California(7). |
| | *Ursus americanus* | American Black Bear | 74(4) | | | 21 | | | |

| Pair | Species (Binomial) | Species (Common) | KT (2n) | % Dist | St.Err | # Seqs | Seq Len | PC | Hybrid Notes |
|---|---|---|---|---|---|---|---|---|---|
| 3 | *Macaca fascicularis* | Crab-eating Macaque | 42(8) | 11.15 | 2.29E-04 | 108 | 1141 | 2 | In 1882, at London Zoo, two F1 hybrids were born (5). Reciprocal crosses are possible, as reported by Chiarelli. One hybrid survived for four months and another for four and a half years(6). At the Yerkes Regional Research Center Field Station, Bernstein reported three captive-born hybrids (one male and two female) that were viable(9). The eldest female was twice backcrossed to *M. fascicularis,* though both infants died(6). An F1 male (*M. fascicularis* x *M. nemestrina*) sired a viable F2 male with a F1 female *M. mulatta?* x *M. nemestrina* (the same male sired F2 viable female offspring with an F1 female *M. nemestrina* x *M. nigra*)(9). This is not a true reciprocal F2 hybrid, there is currently no evidence of breeding experiments using only *M. mulatta* x *M. nemestrina* F1 males and females. If male F1s and female F1s from this pairing are capable of producing F2 offspring, then this could be considered a Category 1 hybrid. |
| | *Macaca nemestrina* | Southern Pig-tailed Macaque | 42(10) | | | 108 | | | |
| 4 | *Macaca mulatta* | Rhesus Macaque | 42(11) | 10.36 | 3.95E-04 | 108 | 1141 | 2 | A female *M. mulatta* x *M. nemestrima* hybrid and male *M. fasicularis* x *M. nemestrima* produced a viable F2 female(9). This is not a true reciprocal F2 hybrid, there is currently no evidence of breeding experiments using only *M. mulatta* x *M. nemestrina* F1 males and females. |
| | *Macaca nemestrina* | Southern Pig-tailed Macaque | 42(10) | | | 108 | | | |

| Pair | Species (Binomial) | Species (Common) | KT (2n) | % Dist | St.Err | # Seqs | Seq Len | PC | Hybrid Notes |
|---|---|---|---|---|---|---|---|---|---|
| 5 | *Papio anubis* | Olive Baboon | 42(12) | 10.17 | 3.42E-04 | 40 | 1141 | 2 | In the Bole Valley in Ethiopia, one adult female and two juvenile males were suspected hybrids of these two species. This observation was strengthened through observations of copulations between the parent species on multiple occasions(13). Two F1 hybrid males are described as fertile, F1 female of this cross was untested. One of the F1 males (Sputnik) was successfully backcrossed with two female *P. hymadryas* baboons and produced 5 viable offspring. The potential for F2 hybrids between F1 hybrids of this pairing were not tested during these breeding experiments. However, when the F1 sputnik was mated with two F1 *P. hamadryas* x *T. gelada* females, 2 of 17 pregnancies resulted in viable offspring with no signs of heterosis (hybrid vigor)(14, 15) |
| | *Theropithecus gelada* | Gelada | 42(14) | | | 10 | | | |
| 6 | *Lepus europaeus* | European Hare | 48(16) | 9.80 | 1.46E-04 | 39 | 1140 | 2 | There is evidence of introgression in Swedish wild individuals, hybrids are therefore at least partially fertile, and can be fully fertile if the mother is *L. timidus* (17-19). A male F1 hybrid produced experimentally had normal testes and motile sperm and was presumed fertile. However, no histology or breeding experiments were performed(18). One study was conducted using breeding experiments to produce F1 hybrids and showed that males had normal sperm and testes(17). |
| | *Lepus timidus* | Mountain Hare | 48(16) | | | 39 | | | |

| Pair | Species (Binomial) | Species (Common) | KT (2n) | % Dist | St.Err | # Seqs | Seq Len | PC | Hybrid Notes |
|---|---|---|---|---|---|---|---|---|---|
| 7 | *Macaca fascicularis* | Crab-eating Macaque | 42(8) | 8.89 | 5.23E-04 | 108 | 1141 | 2 | In Kam Shan Park, Kowloon, China. *M. fasicularis* and *M. thibetana* females care for each other's young(20), and *M. arctoides* could have arisen as a hybrid of the ancient predecessors of *M. thibetana* and *M. fascicularis(21, 22)*. |
| | *Macaca thibetana* | Tibetan Macaque | 42(23) | | | 108 | | | |
| 8 | *Mus musculus domesticus* | Domestic House Mouse | 40(24) | 8.66 | 1.15E-04 | 4 | 1141 | 2 | These species are partially sympatric in Africa and Europe. Both male and some female hybrids are infertile depending on the direction of the cross. Introgression has transferred an anticoagulant rodent poison resistance gene from *M. spretus* into *M. domesticus* in the lab and into wild populations in Germany and Spain(25). F1 males are infertile. F1 females can be backcrossed to male *M.spretus* males but have reduced reproductive capacity. Five out of sixteen male backcross hybrids were fertile, and the remaining nine were infertile. Fertility was confirmed by breeding the males with backcrossed females but not other F1 females(26). |
| | *Mus spretus* | Algerian Mouse | 40(24) | | | 4 | | | |
| 9 | *Mustela erminea* | Stoat | 44(27) | 8.32 | 3.34E-05 | 219 | 1140 | 2 | Listed as “presumed” fertile F1 female and male hybrids, but the reference for this is not listed(6). No direct evidence or breeding experiments to confirm fertility of F1 hybrids. |
| | *Mustela_putorius* | European Polecat | 40(28) | | | 9 | | | |
| 10 | *Ceratotherium simum simum* | Southern White Rhino | 82(29) | 8.24 | 4.29E-03 | 3 | 1140 | 2 | In an 800 hectare enclosure in South Africa's National Zoological Garden's Game Breeding Centre, a female calf was born to a white rhino cow and black rhino bull. Genetic analysis revealed a karyotype of 2n = 83, and the hybrid was subsequently culled(30). |
| | *Diceros bicornis* | Black Rhino | 82(29) | | | 3 | | | |

| Pair | Species (Binomial) | Species (Common) | KT (2n) | % Dist | St.Err | # Seqs | Seq Len | PC | Hybrid Notes |
|---|---|---|---|---|---|---|---|---|---|
| 11 | *Macaca fascicularis* | Crab-eating Macaque | 42(8) | 8.18 | 4.47E-04 | 108 | 1141 | 2 | Numerous suspect hybrids have been observed in the wild in a hybrid zone, and 27 out of 37 individuals in one area were indeterminate in terms of tail length and pelage colouration which suggests hybridisation(31). There is evidence of ancient introgression from *M. mulatta* into *M. fascicularis* with up to 30% of the genome originating from the former(32). Hybridisation also occurs today between both species in a wide hybrid zone running from Vietnam, through Laos, Thailand and probably into Myanmar(33, 34). In 1882, at London Zoo, three F1 hybrids were born(5). A hybrid in Hanover Zoo gave birth to a backcross sired by her father, however, the infant subsequently died(6). |
| | *Macaca mulatta* | Rhesus Macaque | 42(11) | | | 108 | | | |
| 12 | *Acomys dimidiatus* | Eastern spiny mouse | 36-38(35) | 7.96 | 2.92E-04 | 9 | 1144 | 2 | F1 males are sterile(6), but there are no additional references to breeding experiments. |
| | *Acomys minous* | Crete spiny mouse | 38-42(36) | | | 4 | | | |
| 13 | *Cavia aperea* | Brazilian guinea pig | 64(37) | 7.86 | 1.85E-04 | 22 | 1140 | 2 | These two species are able to form fertile hybrids(38), and possibly F2 hybrids(6) since F2 hybrids have been referred to(39). |
| | *Cavia porcellus* | Guinea pig | 64(37) | | | 69 | | | |
| 14 | *Loxodonta africana* | African Bush Elephant | 56(40) | 7.01 | 8.77E-05 | 63 | 1140 | 1 | In Chester Zoo, in 1978 a male calf was born to an Asian cow and an African bull. The infant died 12 days after birth due to necrotising enterocolitis and blood poisoning. This is the only confirmed case of the two species hybridising (41). |
| | *Elephas maximus* | Asian Elephant | 56(40) | | | 9 | | | |

| Pair | Species (Binomial) | Species (Common) | KT (2n) | % Dist | St.Err | # Seqs | Seq Len | PC | Hybrid Notes |
|---|---|---|---|---|---|---|---|---|---|
| 15 | *Canis aureus* | Golden Jackal | 78(42) | 6.16 | 2.01E-03 | 2 | 1140 | 1 | In Northern Africa, jackals have been shown to possess mtDNA lineages from *C. lupus lupaster(43)*. In Bulgaria, nuclear and mitochondrial DNA evidence suggests that *C. lupus* have introgressed with local *C. aureus*. Three individuals are definitely Jackal in origin with regards to mtDNA(44). In captivity, *C. familiaris* x *C. aureus* reciprocal crosses are possible and hybrids are fertile in both sexes(6), though there is no documented captive breeding or evidence of fertility with *C. lupus*. |
| | *Canis lupus* | Wolf | 78(42) | | | 4 | | | |
| 17 | *Loxodonta africana* | African Bush Elephant | 56(40) | 4.57 | 3.42E-05 | 108 | 1140 | 1 | In Africa, a hybrid zone exists around areas of tropical forest. There is evidence of bi-directional introgression with a sex bias of gene flow from larger Bush males to Forest populations. Size-mediated competition between bulls is thought to be the cause. Hybrids are probably fully fertile because there are male Bush elephants with Y-chromosome sequences which cluster with Forest elephant haplotypes, and Bush elephant mtDNA sequences which cluster with Forest elephant mtDNA sequences(45). |
| | *Loxodonta cyclotis* | African Forest Elephant | 56(40) | | | 44 | | | |

| Pair | Species (Binomial) | Species (Common) | KT (2n) | % Dist | St.Err | # Seqs | Seq Len | PC | Hybrid Notes |
|---|---|---|---|---|---|---|---|---|---|
| 18 | *Pan paniscus* | Bonobo | 48(4) | 4.56 | 9.24E-05 | 39 | 1140 | 1 | A small group of chimp/bonobo hybrids was studied (two males and two females) that displayed a mixed array of phenotypes typical of both species with regards to physiology and behaviour(46). A study of three individual bonobos showed no evidence of ancient admixture between chimp subspecies in the wild(47). Fertility of the hybrids is unknown. However, a study analysed the PRDM9 gene in all chimp subspecies(48) revealing that bonobos and Eastern Chimps share an allele for this gene. This is significant since the gene has been known to be a factor contributing to hybrid sterility in mice(49). Asymmetry in this gene can cause failed meiosis during gametogenesis, and because bonobos still share an ancestral allele with chimps, this potentially means hybrid fertility cannot be ruled out. |
| | *Pan troglodytes* | Chimpanzee | 48(4) | | | 40 | | | |
| 20 | *Gorilla beringei graueri* | Eastern Gorilla | 48(4) | 4.39 | 1.85E-04 | 2 | 1140 | 1 | In Central Africa, there is evidence of ancient introgression up to 80kya, and more recent introgression may have also occurred(50). |
| | *Gorilla gorilla gorilla* | Western Gorilla | 48(4) | | | 5 | | | |
| 21 | *Connochaetes taurinus* | Blue Wildebeest | 58(4) | 2.76 | 1.42E-03 | 5 | 1140 | 1 | A population of hybrids was established from two male Blue Wildebeest and 10 Black Wildebeest in 1980. In 1988, no pure animals were observed from 46 air sightings and 15 ground sightings of individuals and groups. During observation, four neonates and two yearlings were observed with hybrid females. This was interpreted as strong circumstantial evidence of F1 hybrid fertility(51). |
| | *Connochaetes gnou* | Black Wildebeest | 58(4) | | | 4 | | | |

| Pair | Species (Binomial) | Species (Common) | KT (2n) | % Dist | St.Err | # Seqs | Seq Len | PC | Hybrid Notes |
|---|---|---|---|---|---|---|---|---|---|
| 22 | *Ceratotherium simum cottoni* | Northern White Rhino | 82(52) | 0.88 | 0.00E+00 | 1 | 1140 | 1 | In captivity, at Dvur Kralove Zoo Czech Republic. A female hybrid was born to a Northern cow and a Southern bull in 1977. Nasi died in 2007(53). Nasi never bred and showed no interest in sex, there is no evidence that oocytes never formed or that oestrus never occurred(52, 54). |
| | *Ceratotherium simum simum* | Southern White Rhino | 82(52) | | | 3 | | | |

1. Lavrov LS & Orlov VN (1973) Karyotypes and taxonomy of modern beavers (Castor, Castoridae, Mammalia). *Zoologische Zhurnal* 52:734-742.
2. Lavrov VL (1996) Hybridisation between East European and Canadian Beavers. *Byulletin Moskovskovo Obschestva*.
3. Kautenburger R, Sander C, Kautenburger R, Sander AC, & Sander K (2008) Population genetic structure in natural and reintroduced beaver (Castor fiber) populations in Central Europe. *25 Animal Biodiversity and Conservation* 312.
4. Kaplan AR (1971) Atlas of Mammalian Chromosomes - Hsu,Tc and Benirschke,K. *Social Biology* 18:325.
5. Flower SS (1928) List of the Vertebrated Aminals Exhibited in the Gardens of the Zoological Society of London 1828-1927.22-25,150-151.
6. Gray AP (1972) Mammalian hybrids. A check-list with bibliography. [Second edition]. *Commonwealth Agricultural Bureaux Slough i-x*:1-262.
7. Hunt B (2012) "Ben the Bear" - A Personal Account.
8. Dutrillaux B, Biemont MC, Viegas-Pequignot E, & Laurent C (1979) Comparison of the karyotypes of four Cercopithecoidae: Papio papio, P. anubis, Macaca mulatta, and M. fascicularis. *Cytogenetics and cell genetics* 23:77-83.
9. Bernstein IS (1974) Birth of two second generation hybrid macaques. *Journal of Human Evolution* 3:205-206.
10. Darlington CD & Haque a (1955) Chromosomes of monkeys and men. *Nature* 175:32.
11. Moore CM*, et al.* (1999) Cytogenetic and fertility studies of a rheboon, rhesus macaque (Macaca mulatta) x baboon (Papio hamadryas) cross: Further support for a single karyotype nomenclature. *American Journal of Physical Anthropology* 110:119-127.
12. Chiarelli B, Koen AL, & Ardito G (1979) Comparative karyology of primates.
13. Dunbar RIM & Dunbar P (1974) On hybridization between Theropithecus gelada and Papio anubis in the wild. *Journal of Human Evolution* 3:187-192.
14. Markarjan DS, Isakov EP, & Kondakov GI (1974) Intergeneric hybrids of the lower (42-chromosome) monkey species of the Sukhumi monkey colony. *Journal of Human Evolution* 3:247-255.
15. Jolly CJ, Woolley-Barker T, Beyene S, Disotell TR, & Phillips-Conroy JE (1997) Intergeneric Hybrid Baboons. *International Journal of Primatology* 18:597-627.
16. Chapman Ja & Flux JEC (1990) Rabbits, hares and pikas: status survey and conservation action plan.61-94.
17. Gustavsson I & Sundt CO (1965) Anwendung von k??nstlicher Befruchtung bei der Hybridisierung von zwei Hasenarten. *Zeitschrift f??r Jagdwissenschaft* 11:155-158.
18. Schröder J, Soveri T, Suomalainen Ha, Lindberg La, & van der Loo W (1987) Hybrids between Lepus timidus and Lepus europaeus are rare although fertile. *Hereditas* 107:185-189.
19. Thulin C-GC, Tegelström H, & Tegelstrm H (2002) Biased geographical distribution of mitochondrial DNA that passed the species barrier from mountain hares to brown hares (genus Lepus): an effect of genetic incompatibility and mating behaviour? *Journal of Zoology* 258:299-306.
20. Burton FD & Chan L (1987) KW. 1987. Notes on the care of long-tail macaque (Macaca fascicularis) infants by stump-tail macaques (Macaca thibetana). *Can. J. Zool* 65:752-755.
21. Tosi aJ, Morales JC, & Melnick DJ (2000) Comparison of Y chromosome and mtDNA phylogenies leads to unique inferences of macaque evolutionary history. *Molecular phylogenetics and evolution* 17:133-144.
22. Tosi AJ, Morales JC, & Melnick DJ (2003) Paternal, maternal, and biparental molecular markers provide unique windows onto the evolutionary history of macaque monkeys. *Evolution; international journal of organic evolution* 57:1419-1435.

23. Fooden J (1986) Taxonomy and evolution of the Sinica group of macaques. 5, Overview of natural history / Jack Fooden. n.s. no.29:40.
24. Silver LM (1995) Mouse Genetics. Concepts and Applications.376.
25. Song Y*, et al.* (2011) Adaptive introgression of anticoagulant rodent poison resistance by hybridization between old world mice. *Current Biology* 21:1296-1301.
26. Hale DW, Washburn LL, & Eicher EM (1993) Meiotic abnormalities in hybrid mice of the C57BL/6J x Mus spretus cross suggest a cytogenetic basis for Haldane's rule of hybrid sterility. *Cytogenetics and cell genetics* 63:221-234.
27. Mandahl N & Fredga K (1980) A comparative chromosome study by means of G-, C-, and NOR-bandings of the weasel, the pygmy weasel and the stoat (Mustela, Carnivora, Mammalia). *Hereditas* 93:75-83.
28. Omodeo P & Renzoni A (1966) The Karyotype of some Mustelidae. *Caryologia* 19:219-226.
29. Trifonov V, Yang F, Ferguson-Smith Ma, & Robinson TJ (2003) Cross-species chromosome painting in the Perissodactyla: delimitation of homologous regions in Burchell’s zebra <i>(Equus burchellii)</i> and the white <i>(Ceratotherium simum)</i> and black rhinoceros <i>(Diceros bicornis)</i>. *Cytogenetic and Genome Research* 103:104-110.
30. Robinson TJ*, et al.* (2005) Interspeci c hybridisation in rhinoceroses: Con rmation of a Black · White rhinoceros hybrid by karyotype, uorescence in situ hybridisation (FISH) and microsatellite analysis. *Society* 6:141-145.
31. Sokrith H, Naven H, & Rawson B (2010) A new record of Macaca fascicularis x M. mulatta hybrids in Cambodia. *Cambodian Journal of Natural History* 1:7-11.
32. Yan G*, et al.* (2011) Genome sequencing and comparison of two nonhuman primate animal models, the cynomolgus and Chinese rhesus macaques. *Nature Biotechnology* 29:1019-1023.
33. Fooden J (1997) Tail Length Variation in Macaca fascicularis and M. mulatta. *PRIMATES* 38:221-231.
34. Hamada Y, Urasopon N, Hadi I, & Malaivijitnond S (2006) Body Size and Proportions and Pelage Color of Free-Ranging Macaca mulatta from a Zone of Hybridization in Northeastern Thailand. *International Journal of Primatology* 27.
35. Barome PO, Lymberakis P, Monnerot M, & Gautun JC (2001) Cytochrome b sequences reveal Acomys minous (Rodentia, Muridae) paraphyly and answer the question about the ancestral karyotype of Acomys dimidiatus. *Molecular phylogenetics and evolution* 18:37-46.
36. GIAGIA-ATHANASOPOULOU EB*, et al.* (2011) New data on the evolution of the Cretan spiny mouse, Acomys minous (Rodentia: Murinae), shed light on the phylogenetic relationships in the cahirinus group. *Biological Journal of the Linnean Society* 102:498-509.
37. Gava A, dos Santos MB, & Quintela FM (2011) A new karyotype for Cavia magna (Rodentia: Caviidae) from an estuarine island and C. aperea from adjacent mainland. *Acta Theriologica* 57:9-14.
38. Rood JP & Weir BJ (1970) Reproduction in female wild guinea-pigs. *Journal of reproduction and fertility* 23:393-409.
39. Carter ND (1972) Carbonic anhydrase isozymes in Cavia porcellus, Cavia aperea and their hybrids. *Comparative Biochemistry and Physiology Part B: Comparative Biochemistry* 43:743-744.
40. Houck ML, Kumamoto aT, Gallagher DS, & Benirschke K (2001) Comparative cytogenetics of the African elephant (Loxodonta africana) and Asiatic elephant (Elephas maximus). *Cytogenetics and cell genetics* 93:249-252.
41. Rees PA (2001) A History of the National Elephant Centre, Chester Zoo. *International Zoo News* 48:170-183.
42. Datta M (1979) Evolutionary Significance of the Chromosome Numbers in Mammals: a point of view. in *Comparative Karyology of Primates* (World Anthropology - De Gruyter), p 75.
43. Gaubert P*, et al.* (2012) Reviving the african wolf canis lupus lupaster in north and west africa: A mitochondrial lineage ranging more than 6,000 km wide. *PLoS ONE* 7.
44. Moura AE*, et al.* (2014) Unregulated hunting and genetic recovery from a severe population decline: The cautionary case of Bulgarian wolves. *Conservation Genetics* 15:405-417.
45. Roca AL, Georgiadis N, & O'Brien SJ (2007) Cyto-nuclear genomic dissociation and the African elephant species question. *Quaternary International* 169-170:4-16.
46. Vervaecke H & Van Elsacker L (1992) Hybrids between common chimpanzees (<i>Pan troglodytes</i>) and pygmy chimpanzees (<i>Pan paniscus</i>) in captivity. *Mammalia* 56:667-669.
47. Prüfer K*, et al.* (2012) The bonobo genome compared with the chimpanzee and human genomes. *Nature* 486:527-531.
48. Groeneveld LF, Atencia R, Garriga RM, & Vigilant L (2012) High diversity at PRDM9 in chimpanzees and bonobos. *PLoS ONE* 7:e39064.

49. Bhattacharyya T*, et al.* (2014) X Chromosome Control of Meiotic Chromosome Synapsis in Mouse Inter-Subspecific Hybrids. *PLoS Genetics* 10:e1004088.
50. Ackermann RR & Bishop JM (2009) Morphological and Molecular Evidence Reveals Recent Hybridisation Between Gorilla Taxa. *Evolution* 64:271-290.
51. Fabricius C, Lowry D, & Van Den Berg P (1987) Fecund black wildebeest x blue wildebeest hybrids. *South African Journal of Wildlife Research* 18:35-37.
52. Houck ML, Ryder Oa, Váhala J, Kock Ra, & Oosterhuis JE (1994) Diploid chromosome number and chromosomal variation in the white rhinoceros (Ceratotherium simum). *The Journal of heredity* 85:30-34.
53. Groves CP, Fernando P, & Robovský J (2010) The sixth rhino: A taxonomic re-assessment of the critically endangered northern white rhinoceros. *PLoS ONE* 5:e9703.
54. Svitalsky M, Vahala J, & Spala P (1993) *Breeding experience with northern white rhinoceros (Ceratotherium simum cottoni) at Zoo Dvur Kralove*).

Supplementary Table 3. A list of hybrid pairs of wild and domestic cats depicted in column B of Figure 1.

| Hybrid Pair | Species (Binomial) | Species (Common) | Karyotype (2n) | % Dist | St.Err | # Seqs | Seq Len | Backcross generations required for fertility |
|---|---|---|---|---|---|---|---|---|
| 1 | *Leptailurus serval* | Serval | 38(1) | 11.28 | 7.51E-04 | 1 | 1137 | The F4 backcross is the first generation in which male fertility can be re-established(2) |
| | *Felis catus* | Cat | 38(3) | | | 4 | | |
| 2 | *Prionailurus bengalensis* | Leopard Cat | 38(4) | 10.94 | 1.06E-03 | 8 | 1137 | The F4 backcross is the first generation in which male fertility can be re-established(2) |
| | *Felis catus* | Cat | 38(3) | | | 4 | | |
| 3 | *Felis chaus* | Jungle Cat | 38(5) | 7.54 | 7.51E-04 | 1 | 1137 | The F4 backcross is the first generation in which male fertility can be re-established(2) |
| | *Felis catus* | Cat | 38(3) | | | 4 | | |

**References**

1. Wurster-Hill DH & Centerwall WR (1982) The interrelationships of chromosome banding patterns in canids, mustelids, hyena, and felids. *Cytogenetics and cell genetics* 34:178-192.
2. Davis BW*, et al.* (2015) Mechanisms Underlying Mammalian Hybrid Sterility in Two Feline Interspecies Models. *Molecular Biology and Evolution*.
3. Menotti-Raymond M*, et al.* (1999) A genetic linkage map of microsatellites in the domestic cat (Felis catus). *Genomics* 57:9-23.
4. Keawmad P, Tanomtong A, & Khunsook S (2007) A Study on Karyotype of the Asian Leopard Cat, Prionailurus bengalensis (Carnivora, Felidae) by Conventional Staining, G-banding and High-resolution Technique. *CYTOLOGIA* 72:101-110.
5. Tanomtong A, Khunsook S, Keawmad P, & Siripiyasing P (2008) Karyological Study of the Jungle Cat, Felis chaus (Carnivora, Felidae) by Conventional Staining, G-banding and High-resolution Staining Technique. *CYTOLOGIA* 73:61-70.

**Supplementary Table 4. NCBI/ENA Accession Numbers for Sequences Used in Distance Calculations**

| Species (Binomial) | NCBI accession numbers (*CYTB*) |
|---|---|
| *Acomys dimidiatus* | AJ010554; AJ010555; AJ233959; AJ233958; AJ012019; Z96061; Z96060; Z96062; AJ012018 |
| *Acomys minous* | AJ233955; AJ233954; AJ233951; AJ233956 |
| *Babyrousa celenbensis* | AJ314559; AY534302; AJ314560; GQ338970; Z50106 |
| *Canis aureus* | AY291433; AQ56603 |
| *Canis latrans* | EU789789; NC_008093; KF661096; DQ480511; DQ480510; DQ480509 |
| *Canis lupus* | AY928668; AY170103; EF689057; AY598499 |
| *Canis lupus lupaster* | HQ845258; EA49881 |
| *Castor canadensis* | FR691684 |
| *Castor fiber* | NC_028625; Q088704; AJ389529; FR691686; FR691687; FR691685; DQ088706; DQ088707; DQ088705; DQ088708; FR691688 |
| *Cavia aperea* | GU136749; GU136748; GU136741; GU136740; GU136743; GU136742; GU136744; GU136747; GU136746; AY382791; AY382790; GU136756; GU136757; GU136754; GU136755; GU136752; GU136753; GU136750; GU136751; GU136758; GU136759; EU544669; GU067538 |
| *Cavia fulgida* | GU136737 |
| *Cavia porcellus* | HM447174; HM447147; HM447146; DQ017044; HM447149; HM447148; DQ017046; HM447170; AY247008; HM447172; GU136732; GU136733; HM447176; NC_000884; AY228363; AY228362; AY228361; HM447171; DQ017042; AF490405; AY382793; HM447181; HM447180; HM447183; HM447182; HM447185; HM447184; HM447187; HM447186; HM447178; HM447179; DQ017045; HM447175; DQ017047; HM447177; DQ017041; DQ017040; DQ017043; HM447173; HM447167; HM447169; HM447168; HM447166; HM447165; HM447164; HM447163; HM447162; HM447161; HM447160; DQ017038; DQ017039; AY245096; AY245097; AY245098; DQ017037; AY245094; AY245095; AJ222767; HM447152; HM447153; HM447150; HM447151; HM447156; HM447157; HM447154; HM447155; HM447158; HM447159 |
| *Ceratotherium simum* | JF718874; Y07726; FJ619038 |
| *Ceratotherium simum cottoni* | FJ619039 |
| *Connochaetes gnou* | AF016637; NC020698; JF728762; JN632626 |
| *Connochaetes taurinus* | JN632628; JN632627; AF016638; NC020699; AF034969 |
| *Dicerorhinus sumatrensis* | FJ905816; AJ245723 |
| *Diceros bicornis* | FJ905814; X56283; EU107377 |
| *Elephas maximus* | D50846; D50844; AY769976; AY769972; AY769973; AY769974; AY769975; AY769977; AB002412 |
| *Equus asinus* | FJ428521; FJ428520; FJ428523; FJ428522; FJ428525; FJ428524; FJ428527; FJ428526; FJ428503; FJ428502; FJ428501; FJ428500; FJ428507; FJ428509; FJ428505; FJ428504; FJ428508; FJ428506; FJ428499; FJ428498; FJ428392; FJ428393; FJ428390; FJ428391; FJ428497; FJ428518; FJ428519; FJ428496; FJ428510; FJ428511; FJ428512; FJ428513; FJ428514; FJ428515; FJ428516; FJ428517; FJ428387; FJ428386; FJ428389; FJ428388 |
| *Equus caballus* | JF511441; JF511449; JF511443; JF511444; JF511447; JF511446; D82932; JF511429; JF511426; JF511432; JF511438; JF511439; JF511428; JF511425; JF511433; JF511434; JF511435; JF511436; JF511437; JF511430; JF511431; JF511458; JF511459; JF511456; JF511457; JF511454; JF511455; JF511452; JF511453; JF511450; JF511451; JF511424; JF511427; JF511440; JF511448; D32190; JF511442; JF511445; JF511423; JF511422 |
| *Felis catus* | AB194814; NC_001700; AB004237; X82296 |
| *Felis chaus* | NC_028307 |
| *Gorilla beringei graueri* | KM242275; KF914213 |
| *Gorilla gorilla gorilla* | NC_011120; D38114; KF914214; NC_001645; EU095336 |
| *Homo sapiens spp. Sima-de-los-Huesos* | KF683087; NC_023100 |
| *Homo sapiens sapiens* (modern) | AF381986; AF381987; AF381984; AF381985; AF381982; AF381983; AF381981; AF382002; AF382003; AF382000; AF382001; AF382006; AF382007; AF381988; AF381989; AF382011; AF382010; AF382013; AF382012; AF382008; AF382009; AF381999; AF381998; AF381991; AF381990; AF381993; AF381992; AF381995; AF381994; AF381997; AF381996; AF382004; KJ669158; AF382005 |

| | |
|---|---|
| *Homo sapiens sapiens* (ancient) | PRJEB6622; KC521457; KC521458; FN600416; KC417443; KU659023; KX638446; KP718913 |
| *Homo sapiens neanderthalensis* | NC_011137; KY751400; KF982693; KU131206; KC879692; KJ533545; KJ533544; FM865407; AM948965; FM865408; FM865409; FM865411; FM865410; KX198088; KX198087; KX198086; KX198085; KX198084; KX198083; KX198082 |
| *Homo sapiens* Denisova | NC_013993; KT780370; KX663333; FR695060 |
| *Leptailurus serval* | NC_028316 |
| *Lepus europaeus* | AY745112; HQ596474; AY745113; HQ596473 |
| *Lepus timidus* | AB687503; AB687502; AB687501; AB687500; AB687507; AB687506; AB687505; AB687504; HM233012; AB687509; AB687508; HQ596483; HM233015; HQ596482; AY599076; AY599075; HM233009; HQ596484; HM232998; AB687530; HM232999; HM232997; AJ279424; AY745109; HM233005; HM232989; HM232988; HM233004; AB687499; AY745108; AY745107; AB687510; AB687511; AB687512; AB687513; AB687514; AB687515; AB687516; AB687517; AB687518; AB687519; HM233007; HM233006; HM233001; HM233000; HM233003; HM233002; HM232990; AY292728; AB058607; AY745105; HM232987; AB687529; AB687528; AY745104; AB687521; AB687520; AB687523; AB687522; AB687525; AB687524; AB687527; AB687526 |
| *Loxodonta africana* | AY768877; AY768849; AY741078; AY768920; AY768921; AY768927; AY768924; AY768925; AY768841; AY768840; AY741073; AY741072; AY768845; AY768844; AY768847; AY741076; AY768934; AY768937; AY768936; AY768838; AY768839; AY768933; AY768932; AY768834; AY768835; AY768836; AY768837; AY768831; AY768832; AY768833; AY768899; AY768892; AY768893; AY768891; AY768897; AY768894; AY768895; AY768902; AY768922; AY768903; AY742801; AY742800; AY768931; AY768930; AY768889; AY768888; AY768885; AY768884; AY768887; AY768886; AY768880; AY741322; AY768938; AY768870; AY768871; AY768872; AY768873; AY768874; AY768875; AY768879; AY768926; AY768848; AY741325; AY741324; AY741326; AY741321; AY741320; AY741323; AY768863; AY768861; AY768860; AY768867; AY768866; AY768865; AY768864; AY768869; AY768868; AY741071; AY768858; AY741070; AY768843; AY768842; AY741075; AY741074; AY741077; AY768846; AY741067; AY768854; AY768852; AY768850; AY768851; AY741068; AY741069; AY768904; AY768905; AY768906; AY768907; AY768900; AY768914; AY768929; AY768919; AY768918; AY768917; AY768916; AY768915; AY768913; AY768912; AY768911 |
| *Loxodonta cyclotis* | AY741080; AY741081; AY768909; AY742799; AY768923; AY768935; AY768898; AY768890; AY768896; AY742802; AY768881; AY768883; AY768882; AY768876; AY768857; AY768878; AY741079; AY741329; AY741328; AY741327; AY768862; AY768859; AY768856; AY768855; AY768853; AY359276; AY359277; AY359274; AY359275; AY359272; AY359273; AY359270; AY359271; AY768901; AY359278; AY359279; AY768928; AY359269; AY359268; AY359265; AY359267; AY359266; AY768908; AY768910 |
| *Macaca arctoides* | KJ567055; AY738634; KM360179; NC_025201 |
| *Macaca fascicularis* | FJ906803; KM851031; KM850998; KJ567052; AF295584; KM851028; KM851029; KM851020; KM851021; KM851022; KM851024; KM851025; KM851026; KM851027; KM851033; KM851032; KM850999; KM851030; KM851037; KM851036; KM851035; KM851034; KM851023; KM851008; KM851009; KM851006; KM851007; KM851004; KM851005; KM851002; KM851003; KM851000; KM851001; KF305937; KM851011; KM851010; KM851013; KM851012; KM851015; KM851014; KM851017; KM851019; KM851018; NC_012670; KM851016 |
| *Macaca mulatta* | KJ567051; KJ567053; AY612638; NC_005943; U38272; JQ821843 |
| *Macaca nemestrina* | HM071128; HM071129; HM071126; HM071127; HM071125; HM071135; HM071134; HM071136; HM071131; HM071130; HM071133; HM071132; EU204975 |
| *Macaca thibetana* | EU294187; KJ567056; AY563620; NC_011519 |

| | |
|---|---|
| *Mus musculus domesticus* | EF108344; FJ374654; FJ374639; FJ374640; FJ374641; FJ374642; FJ374643; FJ374644; FJ374645; FJ374646; FJ374647; FJ374648; FJ374649; AB649457; AB649456; AB649455; AB649459; GQ871746; AB649458; FJ374653; FJ374652; FJ374651; FJ374650; AB649468; AB649469; AB649466; AB649467; AB649464; AB649465; AB649462; AB649463; AB649460; AB649461; AB125774; AB649471; AB649473; AB649472; AB649475; AB649474; AB649477; AB649476; AB649479; AB649478; GQ871745; GQ871744; AB649470; AB649484; AB649480; AB649481; AB649482; AB649483 |
| *Mus musculus musculus* | AB649570; AB649543; AB649545; AB649547; AB649546; AB649549; AB649548; AB205275; AB205274; AB205273; DQ874614; EF108343; AB649571; KF781650; KF781651; KF781652; KF781653; KF781654; KF781655; KF781656; AB649579; AB649572; AB649574; AB649575; AB649576; AB649577; KF781649; KF781648; KF781647; KF781646; KF781645; AB649581; AB649580; AB649582; AB649569; AB649568; AB649563; AB649562; AB649561; AB649560; AB649567; AB649566; AB649565; AB649564; KC663621; AB649558; AB649559; AB649556; AB649557; AB649554; AB649555; AB649552; AB649550; AB649551; AB649578 |
| *Mus spretus* | AY224678; AF159398; KM978950; AY057810; JX457725; AB033700; JX457726 |
| *Mustela erminea* | EF089059; EF089058; EF089055; EF089054; EF089057; AF457442; AF457445; AF457444; EF089053; AF457446; EF089114; EF088988; EF089117; EF088979; EF088978; EF089112; EF088989; EF088975; EF088974; EF088977; EF088976; EF088971; EF088970; EF088973; EF088972; EF089023; EF089089; AF057127; AF457441; EF089109; EF089106; EF089108; AF457443; EF088961; EF089056; EF088947; EF089051; EF089050; EF089028; EF089029; EF088982; EF088983; EF088984; EF088985; EF088986; EF088987; EF089020; EF089021; EF089022; EF089052; EF089024; EF089025; EF089026; EF089027; EF089115; EF088943; EF089116; EF689077; EF689079; EF689078; EF089111; EF089132; EF089133; EF089130; EF089131; EF089134; EF089135; EF088993; EF089019; EF089039; EF089038; EF088997; EF088996; EF088995; EF088994; EF089033; EF089032; EF089031; EF089030; EF089037; EF089036; AB051240; EF089101; EF089118; EF089119; EF088954; EF089086; EF089087; EF089084; EF089085; EF089129; EF089128; EF089080; EF089081; EF089125; EF089124; EF089127; EF089126; EF089121; EF089120; EF089123; EF089122; EF089088; EF089008; EF089009; EF089006; EF089007; EF089004; EF089005; EF089002; EF089003; EF089000; EF089001; GQ153572; EF089100; EF089110; EF089082; EF089011; EF089010; EF089013; EF089012; EF089015; EF089014; EF089017; EF089016; EF089018; EF089107; AF271065; AF271064; AF271066; AF271061; AF271060; AF271063; AF271062; EF088939; EF088967; EF088949; EF088965; EF089064; EF089065; EF089066; EF089067; EF089060; EF089061; EF089062; EF089063; EF089113; EF089068; EF089069; EF088944; EF088945; EF088946; AB564120; EF088940; EF088941; EF088942; EF088992; EF088948; EF088991; EF089099; EF089098; EF089083; EF088990; EF089091; EF089090; EF089093; EF089092; EF089095; EF089094; EF089097; EF089096; EF089077; EF089076; EF089075; EF089074; EF089073; EF089072; EF089071; EF089070; EF088999; EF089079; EF089078; EF088957; EF088956; EF088955; EF088998; EF088953; EF088952; EF088951; EF088950; EF088959; EF088958; EF089035; EF089034; EF089048; EF089049; KM091450; EF089042; EF089043; EF089040; EF089041; EF089046; EF089047; EF089044; EF089045; EF088968; EF088969; EF088962; EF088963; EF088960; EF088966; AB026101; EF088964; EF088980; EF088981; EF089105; EF089104; EF089103; EF089102 |
| *Mustela putorius* | EF987746; AF057128; EF689082; EF689083; X94925; NC_020638; HM106318; AB026107; AB026103 |
| *Myodes centralis* | KJ556626; KJ556625; DQ845185 |

| | |
|---|---|
| *Myodes glareolus* | AF159401; KF918859; FJ881399; FJ881398; FJ881393; FJ881392; FJ881391; FJ881390; FJ881397; FJ881396; FJ881395; FJ881394; AF367083; AF367084; JX477301; JX477288; JX477289; FJ881445; FJ881444; FJ881476; FJ881477; FJ881474; FJ881475; FJ881472; FJ881473; FJ881470; FJ881471; FJ881449; FJ881478; FJ881479; NC_024538; JX477287; KM892829; KM892828; KM892825; KM892827; KM892826; KM892821; KM892820; KM892822; JX477273; JX477272; JX477271; JX477270; JX477277; JX477276; JX477275; JX477274; JX477279; JX477278; AF119272; FJ881432; FJ881433; FJ881430; FJ881431; FJ881436; FJ881437; FJ881434; FJ881435; FJ881438; FJ881439; JX477303; JX477302; JX477307; JX477306; JX477305; JX477304; AM392368; KM892814; KM892815; KM892816; KM892817; KM892810; KM892811; KM892812; KM892813; JX477282; JX477283; JX477280; JX477281; KM892818; KM892819; JX477284; JX477285; AF318584; AF318585; FJ881469; FJ881468; FJ881461; FJ881460; FJ881463; FJ881462; FJ881465; FJ881464; FJ881467; FJ881466; FJ881452; FJ881455; FJ881458; FJ881459; KM892841; KM892840; FJ881425; FJ881424; FJ881427; FJ881426; FJ881421; FJ881420; FJ881423; FJ881422; FJ881429; FJ881428; JX477286; JX477309; JX477308; FJ881450; FJ881451; AY309419; FJ881453; FJ881454; FJ881456; FJ881457; JX477300; AF367079; JX477299; JX477298; JX477295; JX477294; JX477297; JX477296; JX477291; JX477290; KM892809; JX477292; FJ881448; FJ881418; FJ881419; FJ881414; FJ881415; FJ881416; FJ881417; FJ881410; FJ881411; FJ881412; FJ881413; FJ881480; FJ881389; JX477293; FJ881443; FJ881442; FJ881441; FJ881440; FJ881447; FJ881446; AY309420; AY309421; KM892838; KM892839; KM892830; KM892831; KM892836; KM892837; KM892834; JX477265; JX477266; JX477267; JX477268; JX477269; FJ881409; FJ881408; FJ881407; FJ881406; FJ881405; FJ881404; FJ881403; FJ881402; FJ881401; FJ881400; JX477310; JX477311; JX477312; JX477313; JX477314; JX477315; JX477316 |

| | |
|---|---|
| *Myodes rutilus* | JF430972; JF430971; KJ789546; KJ789547; KJ789423; KJ789422; KJ789421; KJ789420; KJ789427; KJ789426; KJ789425; KJ789424; JF430970; AB031581; JQ182053; JQ182052; JQ182055; JQ182054; JQ182057; JQ182056; JQ182059; JQ182058; KJ789510; KJ789580; KJ789581; KJ789582; KJ789584; KJ789585; KJ789467; KJ789465; KJ789464; KJ789463; KJ789462; KJ789461; AF119274; KJ789468; AF272631; AF272632; AF272638; KJ789419; HM165375; HM165373; KJ789519; KJ789518; KJ789513; KJ789512; KJ789511; KJ789411; KJ789517; KJ789516; KJ789515; KJ789514; JQ182082; JQ182083; JQ182080; JQ182081; JQ182086; JQ182087; JQ182084; JQ182085; JQ182088; JQ182089; KJ789466; KJ556636; KJ789460; JF430948; JF430949; JF430942; JF430943; JF430940; JF430941; JF430946; JF430947; JF430944; JF430945; AB072207; AB072208; AB072209; JQ182068; JQ182069; JQ182060; JQ182061; JQ182062; JQ182063; JQ182064; JQ182065; JQ182066; JQ182067; KJ789555; KJ789554; KJ789552; KJ789418; KJ789550; KJ789416; KJ789417; KJ789415; KJ789410; KJ789558; JX477341; JX477340; JF430963; JF430959; KJ789593; KJ789458; KJ789459; KJ789597; KJ789595; KJ789594; KJ789452; KJ789453; KJ789450; KJ789451; KJ789456; KJ789457; KJ789454; KJ789429; KJ789428; KJ789528; KJ789526; KJ789527; KJ789524; KJ789525; KJ789522; KJ789523; KJ789520; KJ789521; JX477339; KJ789548; KJ789549; KJ556622; JF430958; JF430955; JF430954; JF430957; JF430951; JF430950; JF430953; JF430952; AB072215; AB072214; AB072217; AB072216; AB072211; AB072210; AB072213; AB072212; KJ789430; AB072219; AB072218; KJ789496; KJ789494; KJ789495; KJ789492; KJ789493; KJ789490; KJ789491; KJ789498; KJ789499; JQ182079; JQ182078; JQ182073; JQ182072; JQ182071; JQ182070; JQ182077; JQ182076; JQ182075; JQ182074; KJ789568; KJ789569; KJ789562; KJ789560; KJ789567; KJ789564; KJ789565; KJ789405; KJ789404; KJ789407; KJ789406; KJ789409; KJ789408; JQ231219; JF430964; JF430965; JF430966; JF430967; JF430960; JF430961; JF430962; JF430968; JF430969; AB072224; AB072220; AB072221; AB072222; AB072223; KJ789449; KJ789448; KJ789445; KJ789444; KJ789447; KJ789446; KJ789441; KJ789440; KJ789443; KJ789442; KJ789531; KJ789530; KJ789533; KJ789532; KJ789535; KJ789534; KJ789537; KJ789539; KJ789538; KJ789433; KJ789434; KJ789435; KJ789436; KJ789437; KJ789438; KJ789439; AY309428; KJ789431; AY309426; AY309427; AY309424; AY309425; KJ789489; KJ789488; KJ789481; KJ789480; KJ789483; KJ789482; KJ789485; KJ789484; KJ789487; KJ789486; JF430939; JF430938; JF430937; JF430936; JF430935; JF430934; KJ789579; KJ789578; KJ789575; KJ789574; KJ789576; KJ789571; KJ789573; KJ789572; KJ789476; KJ789477; KJ789470; KJ789471; KJ789472; KJ789473; KJ789478; KJ789479; KJ789509; KJ789500; KJ789501; KJ789502; KJ789503; KJ789504; KJ789505; KJ789506; KJ789507; KJ556724 |
| *Oryctolagus cuniculus* | AY292717; U07566; HQ596486; KT626640 |
| *Pan paniscus* | GU189668; GU189669; GU189664; GU189665; GU189666; GU189667; GU189660; GU189661; GU189662; GU189663; GU189677; GU189676; GU189675; GU189674; GU189673; GU189672; GU189671; GU189670; D38116; GU189657; GU189659; GU189658; JF727228; JN191197; JN191194; JN191188; JN191189; JF727238; JF727231; JF727232; JN191198; JN191196; JN191195; JN191193; JN191192; JN191191; JN191190; NC_001644; HM015213 |
| *Pan troglodytes* | JN191232; JN191233; JN191230; JN191231; JN191234; JN191235; KP317203; JN191225; JN191224; JN191226; JN191221; JN191220; JN191223; JN191222; JN191229; JN191228; JF727176; JF727173; JF727179; D38113; JF727166; JN191227; JN191215; JN191216; JN191210; JN191211; JN191212; JN191213; JN191218; JN191219; JN191214; JN191217; NC_001643; JN191209; EU095335 |
| *Panthera leo* | GU131184; KC495058; HM107681; NC_028302.1 |
| *Panthera tigris* | AF053040; AF053030; AF053021; KP202268.1 |
| *Papio anubis* | KM267407; JX946198; JX946197; EU885461; EU885460; EU885456; EU885457; EU885454; EU885455; EU885452; EU885453; EU885450; EU885451; EU885458; EU885459 |
| *Papio cynocephalus* | GQ148706; GQ148698; GQ148699; GQ148708; GQ148709; GQ148704; GQ148705; GQ148707; GQ148700; GQ148701; GQ148702; GQ148703; GQ148711; GQ148710; EU885440 |

| | |
|---|---|
| *Papio hamadryas* | KM267345; JX946201; KM267347; KM267343; KM267382; KM267383; KM267380; KM267381; KM267386; KM267387; KM267384; KM267385; KM267388; KM267389; KM267346; KM267379; KM267378; KM267395; KM267394; KM267397; KM267396; KM267391; KM267390; KM267393; KM267392; KM267398; KM267377; KM267376; KM267339; KM267404; KM267401; KM267400; KM267403; KM267402; KM267342; KM267368; KM267369; KM267364; KM267365; KM267366; KM267367; NC_001992; KM267370; KM267373; KM267372; KM267371; KM267375; EU885441; EU885443; EU885442; EU885445; EU885444; EU885446; Y18001; KM267350; KM267348; KM267349; KM267344; KM267399; KM267354; Y16590; KM267374; KM267351; KM267353; KM267352; KM267355; KM267356 |
| *Peromyscus maniculatus* | JF489123; FJ415093; FJ415092; FJ415095; FJ415094 |
| *Peromyscus nasutus* | AF155399; AY376426 |
| *Peromyscus polionotus* | JF322887; JF322886; EU140786; EU140787; EU140783; EU140780; EU140788; EU140789; EU140764; EU140765; EU140766; EU140767; EU140760; EU140761; EU140762; EU140763; EU140768; JF322889; JF322888; EU140791; EU140790; EU140792; EF216339; EU140793; EU140777; EU140776; EU140775; EU140774; EU140773; EU140772; EU140771; EU140770; EF216336; EF216337; EU140779; EU140778; JF322896; JF322897; JF322894; JF322895; JF322892; JF322893; JF322890; EU140759; JF322891; EU140758; EF216338; EU140785; EU140782; EF216347; EF216346; EF216345; EF216344; EF216343; EF216342; EF216341; EF216340; EU140781; EU140757; EU140756; EU140769; EU140784; JF322885 |
| *Peromyscus truei comanche* | AY376428; AY376429; AY376430; AY376431 |
| *Pongo abelii* | X97707; NC_002083; U38274 |
| *Pongo pygmaeus* | D38115; NC_001646; X97717 |
| *Prionailurus bengalensis* | AB194818; AB210231; AB210238; AB210227; NC_028301.1; KP202260.1; KP202259.1; KP202258.1; KP202257.1 |
| *Rhinoceros sondaicus* | FJ905815; AJ245725 |
| *Rhinoceros unicornis* | JF718877; X97336 |
| *Scotinomys teguina* | AF108705; JN851815 |
| *Sus domesticus* | KJ746662; KJ746663; KJ746664; KJ746665; KJ746666; AP003428; KF971862; NC_012095; KC469586; KC469587 |
| *Theropithecus gelada* | JQ257000; FJ785426; KM267405; KC757412; EU885487; NC_019802 |
| *Ursus americanus* | U23556; AF303109; NC_003426; AF268264; AF268263; AF268262; AF268268; X82307; AF268258; AF268259; AF268270; KM257060; AF268271; AF268265; JX196366; AF268267; AF268266; AF268261; AF268260; AF268269; KM257059 |
| *Ursus arctos* | HQ685901; HQ685903; HQ685902; HQ685905; HQ685904; HQ685907; HQ685906; HQ685909; HQ685908; JX196367; HQ685912; GU573491; HQ685928; HQ685927; HQ685930; HQ685931; HQ685932; HQ685933; HQ685926; HQ685923; HQ685922; HQ685921; HQ685920; HQ685925; HQ685924; HQ685918; HQ685919; HQ685916; HQ685917; HQ685915; HQ685913; HQ685910; HQ685911; HQ685929 |
| *Ursus maritimus* | AJ428577; U18898; X82309; AP012597; AP012594; AP012595; GU573490; GU573488; GU573485; JX196392; JX196390; JX196391; JX196374; JX196375; JX196376; JX196377; JX196370; JX196371; JX196372; JX196373; JX196378; JX196379; JX196389; JX196388; JX196381; JX196380; JX196383; JX196382; JX196385; JX196384; JX196387; JX196386 |
| **Species (Binomial)** | **NCBI accession numbers (full mtDNA genome)** |
| *Pan paniscus* | JN191188; JF727223; GU189677; GU189658; GU189670 |
| *Macaca fascicularis* | FJ906803; KM851031; KF305937; KM851007; KM851023 |
| *Papio anubis* | JX946198; JX946197 |
| *Theropithecus gelada* | KC757412; FJ785426 |
| *Pan troglodytes* | JN191217; JN191223; JN191207; JN191203; JN191201 |
| *Macaca nemestrina* | KP765688 |
| *Homo sapiens sapiens* | JX153451; HUMMTCG; HQ610990; HQ610992; HQ610993 |
| *Papio hamadryas* | JX946201; Y18001 |
| *Macaca mulatta* | AY612638; KJ567053; JQ821843 |

| **Species (Binomial)** | **NCBI accession numbers (*CHRNA1*)** |
|---|---|
| *Pan paniscus* | HM763406 |
| *Macaca fascicularis* | HM763362 |
| *Papio anubis* | HM763377 |
| *Theropithecus gelada* | HM763389 |
| *Pan troglodytes* | HM763408; HM763407 |
| *Macaca nemestrina* | HM763366 |
| *Homo sapiens sapiens* | HM763397 |
| *Macaca mulatta* | HM763365 |
| *Papio hamadryas* | HM763378 |
| **Species (Binomial)** | **NCBI accession numbers (*ZFX*)** |
| *Pan paniscus* | HM757090 |
| *Macaca fascicularis* | HM757045 |
| *Papio anubis* | HM757061 |
| *Theropithecus gelada* | HM757072 |
| *Pan troglodytes* | HM757092; HM757091 |
| *Macaca nemestrina* | HM757049 |
| *Homo sapiens sapiens* | HM757080 |
| *Macaca mulatta* | HM757048 |
| *Papio hamadryas* | HM757062 |
| **Species (Binomial)** | **NCBI accession numbers (*ZFY*)** |
| *Pan paniscus* | HM756983 |
| *Macaca fascicularis* | HM756948 |
| *Papio anubis* | HM756960 |
| *Theropithecus gelada* | HM756968 |
| *Pan troglodytes* | HM756984 |
| *Macaca nemestrina* | HM756951 |
| *Homo sapiens sapiens* | HM756974 |
| *Papio hamadryas* | HM756961 |
| *Macaca mulatta* | HM756950 |
| **Species (Binomial)** | **NCBI accession numbers (*GHR*)** |
| *Pan paniscus* | HM761455 |
| *Macaca fascicularis* | HM761390 |
| *Papio anubis* | HM761400 |
| *Theropithecus gelada* | HM761403 |
| *Pan troglodytes* | HM761456; HM761452 |
| *Macaca nemestrina* | HM761439 |
| *Homo sapiens sapiens* | HM761441 |
| *Papio hamadryas* | HM761401 |
| *Macaca mulatta* | HM761424 |

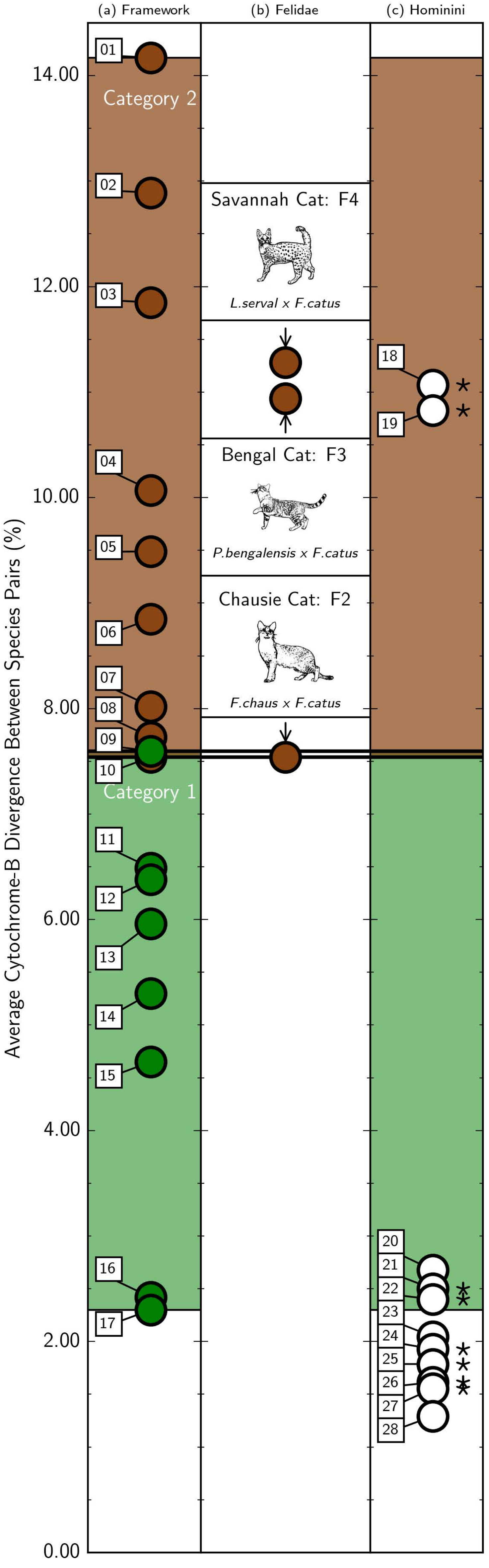
(a) Framework
(b) Felidae
(c) Hominini
Average Cytochrome-B Divergence Between Species Pairs (%)
14.00
12.00
10.00
8.00
6.00
4.00
2.00
0.00
Category 2
Category 1
Savannah Cat: F4
L.serval x F.catus
Bengal Cat: F3
P.bengalensis x F.catus
Chausie Cat: F2
F.chaus x F.catus

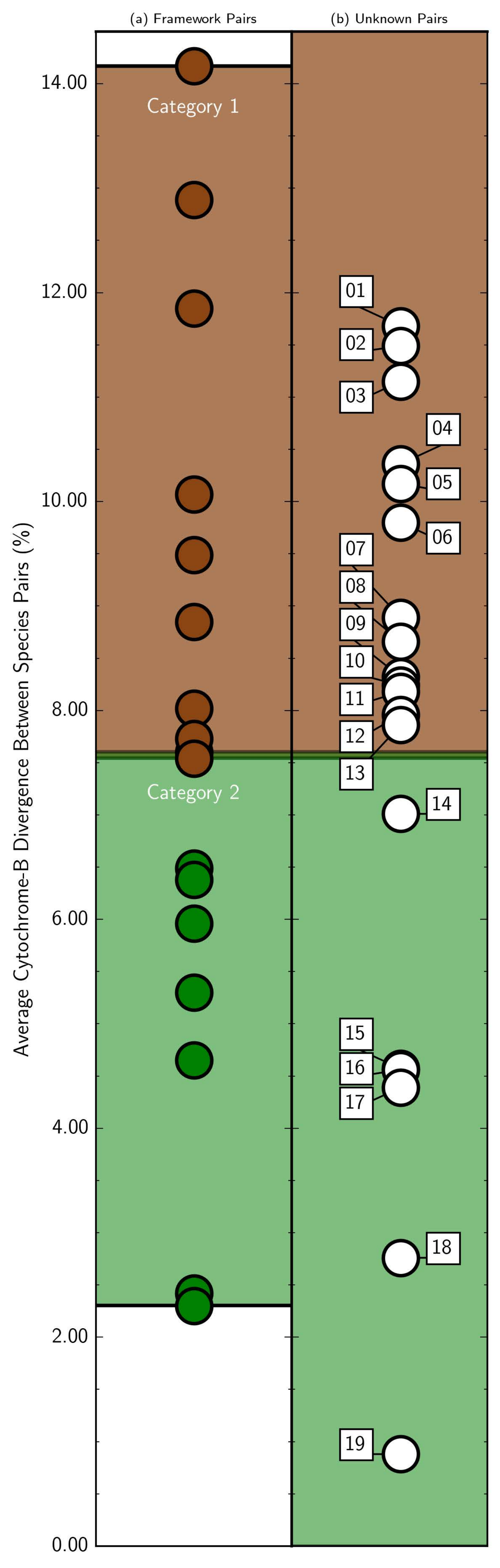
(a) Framework Pairs
(b) Unknown Pairs
Average Cytochrome-B Divergence Between Species Pairs (%)
14.00
12.00
10.00
8.00
6.00
4.00
2.00
0.00
Category 1
Category 2

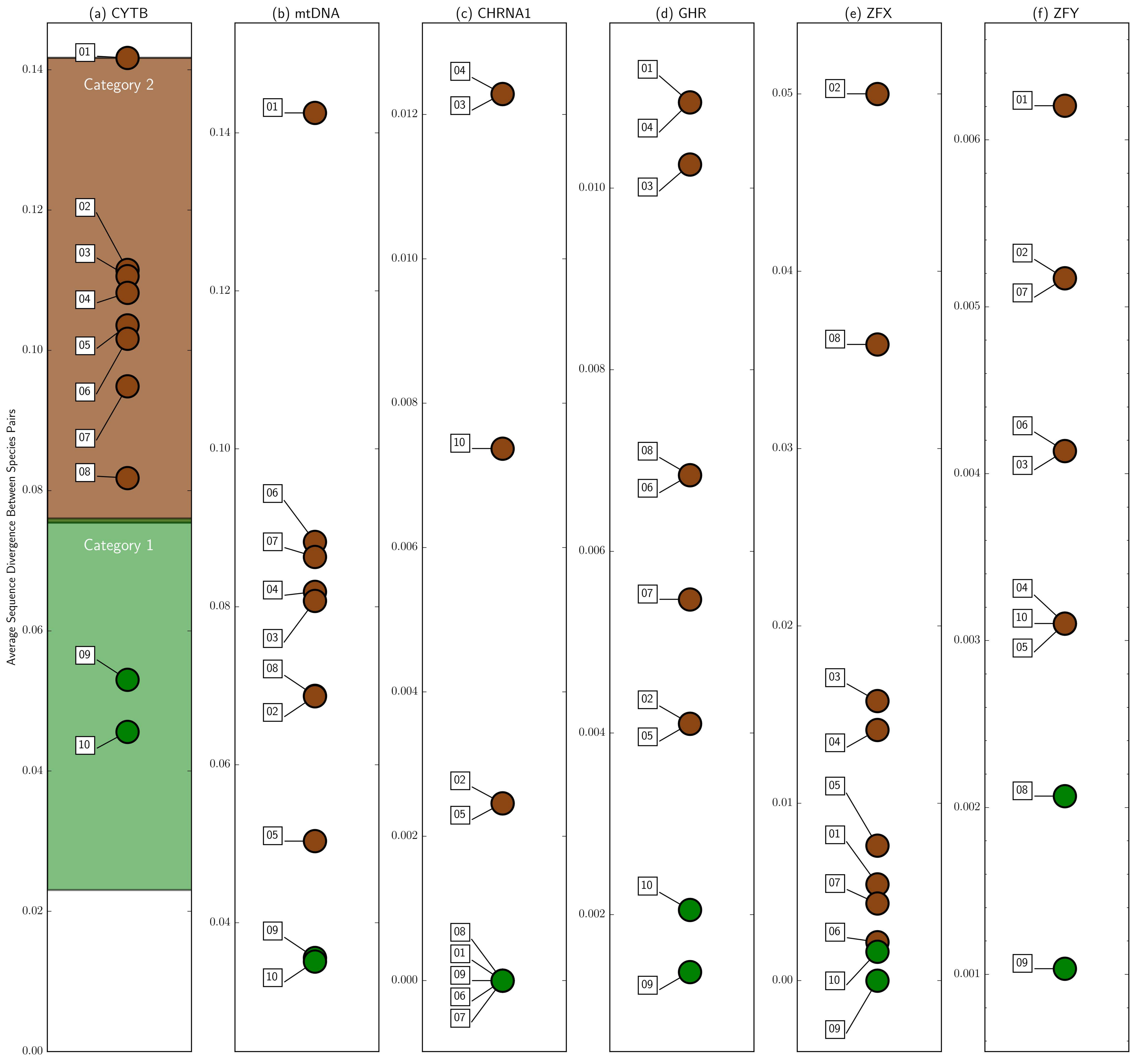

(a) CYTB
(b) mtDNA
(c) CHRNA1
(d) GHR
(e) ZFX
(f) ZFY
Average Sequence Divergence Between Species Pairs
Category 2
Category 1

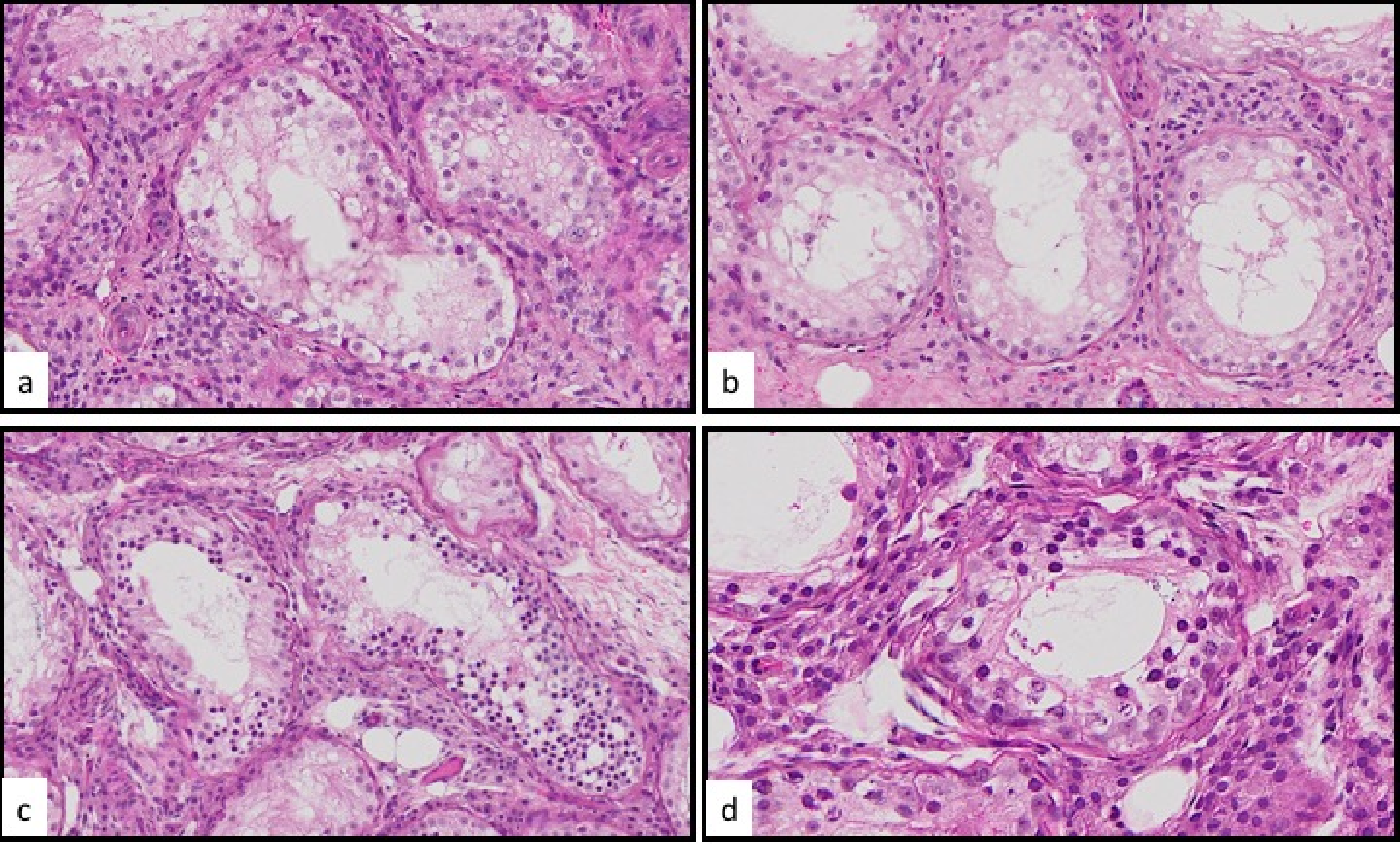
a
b
c
d
